# Antimitotic Activity of *Samanea saman* Leaf Extract in the In Vitro Development of *Tripneustes gratilla* Embryo

**DOI:** 10.1101/2023.05.10.540278

**Authors:** Justin Riley Y. Lam, Guinevere A. Casimpan, Christine May L. Batoy, Ma. Felaine Cebedo, Jann Ycleo T. Cuesta, Jecell Gervacio, Joshua Benjamin R. Grapa, Ma. Katrina Ada F. Mabelin, Merry Grace S. Pepito, Justine Mae A. Rama, Daniel Joseph Z. Simporios, Zackaree Michael A. Villanueva, Jolienne Abbygail M. Villaruel, D. Pepito Gwendolyn, Christopher A. Lu Adrian

**Affiliations:** Cebu Institute of Medicine

**Keywords:** antimitotic activity, acacia, sea urchin, embryo

## Abstract

**TITLE:** Antimitotic Activity of *Samanea saman* Leaf Extract in the In Vitro Development of *Tripneustes gratilla* Embryo

**INTRODUCTION:** Commercially available anticancer drugs are expensive and may have side effects. This led the researchers to look into alternative plant sources. One of these is acacia or *S. saman*.

**OBJECTIVE:** To determine whether *S. saman* leaf extract possesses antimitotic activity on *T. gratilla* embryos using vincristine sulfate as positive control.

**METHODOLOGY:** *S. saman* leaves were extracted using hexane, CCl_4_, CHCl_3_, and water. *T. gratilla* embryos were fertilized in vitro. The fractions, vincristine sulfate, and DMSO in filtered sea water were added with the fertilized embryos in petri dishes. Samples were taken at 15 minutes after fertilization and every 30 minutes thereafter until the negative control reached a 32-cell stage. Fifty cells and their mean cell stage was evaluated per treatment. One-way ANOVA and Tukey tests were performed.

**RESULTS:** Significant differences were seen starting at 75 minutes post fertilization up until the 165^th^ minute using One-way ANOVA. The Tukey test showed that all aqueous and CCl_4_ as well as hexane (200, 400, 800 ppm) extracts had no significant difference compared to vincristine sulfate; while all CHCl_3_, hexane (100 ppm), and CCl_4_ (100 ppm) extracts showed a significant difference compared to vincristine

**CONCLUSION:** The aqueous, hexane, and carbon tetrachloride extracts possess potential antimitotic activity on *T. gratilla* embryos. Thus, it is a potential alternative to vincristine sulfate as an antimitotic agent.

## CHAPTER 1: INTRODUCTION

### Background of the Study

Cancer is a general term for diseases that primarily involve the abnormal proliferation of cells caused by mutations in the DNA sequences that control cell division, and these mutations are generally genetic in nature.^1^ Other factors that predispose an individual to cancer include exposure to carcinogens in the environment, and practice of lifestyle-associated risk factors such as habitual smoking and alcohol consumption.^2^

As of 2018, the World Health Organization (WHO) considers cancer as the second leading cause of death globally, accounting for an estimated 9.6 million deaths in 2018; about 1 out of 6 deaths is due to cancer.^3^ According to the Philippine Statistics Authority, cancer was responsible for around 10.4% deaths in the country in 2016, making it the second leading cause of mortality.^4^ Thus, it is considered one of the foremost concerns in the health sector both locally and internationally.

Treatment modalities of cancer differ depending on its nature and severity.^5^ Common treatments include surgery, radiation, and chemotherapy, which may include the use of antimitotic agents.^6^ Antimitotic agents are substances that block the progression of the cell cycle, inhibiting mitosis, and eventually cause cell death.^5^ The antiproliferative capability of antimitotic agents make it one of the focuses in the development of anticancer drugs.

More than 60% of anticancer agents are formulated from natural sources like plants, marine organisms, or microorganisms. These natural products therefore serve as high-impact sources for novel compounds for potential therapeutic agents. *Paclitaxel*, *combretastatin*, *bryostatin* and *discodermolide* are examples of anticancer drugs derived from natural resources. These highlight the importance of research among natural products in the field of cancer chemotherapy.^7^

A variety of plants have anti-inflammatory, free radical-scavenging, antioxidative, and antimitotic potential capabilities.^8^ Plants produce antioxidants such as flavonoids and polyphenolics to prevent oxidative stress caused by reactive oxygen species. Plants with high concentrations of antioxidants may have a role in the prevention of cancer development.^9^ In several studies, an inverse relationship between dietary intake of foods rich in antioxidants and the incidence of disease have been noted.^10^

The acacia tree (*Samanea saman*), has been shown to have a diverse array of applications as traditional medicine in different cultures. The bark is boiled and used to treat constipation, while the inner bark and leaves are traditionally used to treat diarrhea. In Venezuela, literature has also pointed to the tree’s roots being used to treat stomach cancer.^11^ Extract of its flowers has exhibited some moderate anticancer activity when tested against human breast cancer lines and has shown excellent free radical-scavenging activity.^12, 13^ Preliminary phytochemical screening showed that *S. saman* contains tannins, flavonoids, steroids, saponins, cardiac glycosides, terpenoids, alkaloids, and phenols.^14, 15^

The constant rise of cancer cases in today’s generation with its high morbidity and mortality has challenged the medical community to seek out and research new treatment modalities and new sources of potential therapeutic agents, especially those found in plants used in traditional medicine. These recent treatment modalities may also serve as an answer to cases of malignancy that are unresponsive to present chemotherapy.

### Statement of the Problem

Current anticancer drugs have severe side effects—most notably acquiring resistance and cytotoxic effects of the drugs on non-tumorigenic cells.^6^ Thus, there is a continued search for new compounds that are at par or more effective in arresting mitotic activity, but with minimal side effects. Plants are the main source of these antimitotic agents—vinca alkaloids *vincristine* and *vinblastine*, *paclitaxel* and *etoposide* are some examples of these plant-derived anticancer drugs.^16^ Therefore, looking into plant families with potential medicinal value is an important strategy of developing new possible anticancer agents. Recent studies on *S. saman* proved that it contains antioxidants, which may point to a potential antimitotic capability.

This research may also serve as a baseline study for more comprehensive research on the phytochemical substances present in *S. saman* leaves.

### Research Question

Does *S. saman* leaf extract possess antimitotic activity on *T. gratilla* embryos?

### Research Objectives

The main objective of the study is to determine the antimitotic activity of *S. saman* leaf extract against the in vitro development of *T. gratilla* embryo.

Specifically, the study is designed:

1. To determine the mean cell stages of *T. gratilla* embryo at fixed time intervals of treatment with different concentrations of *S. saman* hexane, carbon tetrachloride, chloroform, and aqueous leaf extract.
2. To determine the mean cell stages of *T. gratilla* embryo at fixed time intervals of treatment with 20 μg/ml vincristine sulfate (positive control) and filtered seawater group in dimethyl sulfoxide solution (negative control).
3. To compare the mean cell stages of *T. gratilla* embryo at fixed time intervals of treatment with different concentrations of *S. saman* hexane, carbon tetrachloride, chloroform, and aqueous leaves extract, against 20 μg/ml vincristine sulfate group and filtered seawater group in dimethyl sulfoxide solution.

### Research Hypothesis

**H_o_:** There is no significant difference in the antimitotic activity between vincristine sulfate and the different *S. saman* hexane, carbon tetrachloride, chloroform, and aqueous leaf extracts.

**H_a_:** There is a significant difference in the antimitotic activity between vincristine sulfate and the different *S. saman* hexane, carbon tetrachloride, chloroform, and aqueous leaf extracts.

## CHAPTER 2: REVIEW OF RELATED LITERATURE

### Samanea saman

*S. saman* is a member of the Fabaceae family cultivated and naturalized throughout the tropics. In the Philippines, it is commonly known as “acacia”, or “palo de China”. It has a characteristic umbrella-shaped canopy that serves as shade on small farms, along roads, parks and pastures in the Pacific.^18^

*S. saman* has a number of environmental uses such as nitrogen enrichment for soil, as shade, windbreaks, silvopasture, as firewood or timber, as fiber for weaving and clothing, as resin or glue, and as food for animals and birds. Some parts of the tree are used for medicinal purposes as well as a remedy for some health problems. In the Philippines, a mixture made of the inner bark and fresh leaves is used for treatment of diarrhea. In Venezuela, the roots are used for stomach cancer where it is applied in a hot bath. In the West Indies, the seeds of the tree are used as a medicinal remedy for sore throat by chewing these seeds.^18^ Because of the multiple traditional uses of this plant, several studies have been conducted to further explore its potential.^8^

The extract of the flowers of the *S. saman* has exhibited some moderate anticancer activity when tested against human breast cancer cell lines.^12^ *S. saman* also shows excellent free radical-scavenging activity and preliminary phytochemical screening has exhibited it to be positive for tannins, flavonoids, steroids, saponins, cardiac glycosides, terpenoids alkaloids and phenols. The ability of a compound to sequester highly reactive free radicals is an important property in cancer treatment as its reactivity to cells usually preludes cancer.^12–14^ It was also noted that synthesis of zinc nanoparticles artificially synthesized from leaf extracts exhibited genotoxic properties by eliciting chromosomal abnormalities on mitotic root meristems, which further proves its antimitotic capability.^12^

### Phytochemical Constituents of *S. saman*

James Kennard S Jacob (2016) determined the phytochemical constituents and the inhibitory activities of *S. saman* pods. Powdered pods were subjected to ethanol and aqueous extraction. Alkaloids, saponins, tannins, glycosides, steroids, terpenoids, and resins were present in both extracts, however, absence of flavonoids in both extraction was observed. *S. saman* pods revealed a significant inhibitory effect against in vitro bioassay of *Fusarium oxysporum*, known to cause wilting on crops, and the two bacterial pathogens *Escherichia coli* and *Staphylococcus aureus* via disc diffusion method.^19^

The bark of *S. saman* was tested for the antiseptic potential of its alkaloids against *E. coli, S. aureus,* and *B. cereus*.^20^ The extraction of the alkaloid was done through continuous extraction method. Two extracts, crude extracts and alkaloid rich fraction, were isolated and partially purified from the bark using toluene:acetone:ethanol:ammonia as solvent in thin layer chromatography. Gentamicin sulfate was used as the reference substance since it exhibits that the 3 microorganisms were all susceptible and exhibited complete inhibition of *E. coli*. Among the three organisms used, only the *S. aureus* plate showed complete inhibition by the two extracts of the bark and showed no zone of inhibition in the other two. It was then concluded that the extracts of the bark can be used as an antibacterial agent and as a source of alternative medicine.^21^

In a study by S. Prema, et. al. (2018), the bark and leaves of the *S. saman* were tested for their phytochemical constituents. Using the Soxhlet apparatus, successive extraction of the powderized bark and leaves was done with the solvent mixture of methanol and 50% hydroethanol. Qualitative analysis of the phytochemical screening was done with methanol and 50% hydroethanol extract. Alkaloids, flavonoids, coumarins, saponins, steroids, tannins, terpenoids, phenols, and cardiac glycosides are secondary metabolites present when screened using standard phytochemical methods such as Dragendroff’s test, Shinoda test, Lead acetate test, Alkaline reagent test, Liebermann-Burchard test, Salkowski test, Ferric chloride test, and Keller-Killiani’s test.^14^

A study conducted by P. Arulpriya, et. al. (2010) on the extract of the fallen parts of the *S. saman* showed that higher antioxidant potential of the extracts was observed in both 1,1-diphenyl-2-picrylhydrazyl (DPPH) radical-scavenging assay and reducing power assay. The petroleum ether, ethyl acetate, chloroform, aqueous and HCl extracts of *S. saman* were examined for their phytochemical constituent and for their in vitro activities. Most of the extracts showed excellent free radical-scavenging activity.^12^ Literature reports provide evidence that the reducing power of bioactive compounds is associated with antioxidant activity.^8,10,12,13^

P. Arulpriya, et. al. (2010) also studied the electrochemical behavior of ethyl acetate extract of *S. saman* using cyclic voltammetry three electrode system. The extracts were investigated for the presence of phytochemical constituents and antioxidant activity assessed from its oxidation potential values at glassy carbon electrode. The results showed that the ethyl acetate (EA) extract exhibited presence of alkaloids, flavonoids, tannins, saponins, quinine, phenol, betacyanin and glycosides.^12^

A. Ravi Kumar (2013) concluded that alkaloids, flavonoids, carbohydrates, glycosides and tannins were present in *S. saman*. The phytochemicals of *S. saman* were evaluated using the crude drug powder extracts of the leaves.^22^

Ferdous, et. al. (1970) investigated the n-hexane, carbon tetrachloride and chloroform soluble fractions of crude methanolic extract of *S. saman* bark. The fractions were tested for antioxidant, antimicrobial, and cytotoxic potential. Antioxidant activity of the extracts was evaluated by DPPH radical-scavenging assay and total antioxidant activity test. Chloroform and hexane soluble fraction showed IC50 value of 12 μg/ml and 14 μg/ml respectively in scavenging DPPH radical while the reference, butylated hydroxytoluene showed an IC50 value of 10 μg/ml. The carbon tetrachloride fraction showed the highest total antioxidant capacity. The carbon tetrachloride fraction was also found to possess mild to moderate microbial growth inhibitory capacity. Antimicrobial activity was tested using disc diffusion method against thirteen bacteria and three fungi and cytotoxicity was tested by brine shrimp lethality bioassay. In the brine shrimp lethality bioassay, the n-hexane, carbon tetrachloride, chloroform soluble fractions showed LC50 value of 14.94 μg/ml, 0.831 μg/ml and 3.288 μg/ml respectively. The results suggest good antioxidant and cytotoxic potential of chloroform and hexane soluble fractions and antimicrobial activity of carbon tetrachloride fraction of *S. saman* bark extract.^23^

### Kupchan Extraction

Liquid–liquid extraction is considered a traditional separation method used to partition analytes based on their solubility in two different liquid solvents.^24^ A well-described methodology was published by Kupchan in 1969 which involved isolation of several tumor inhibitory terpenoids from various plant species. His methodology allows separation between the polar and the non-polar compounds, incorporating systemic fractionation, guided by biological assays allowing minor constituents to be isolated.^25^ VanWegenen, et. al. (1993) provided a modified version of the Kupchan method.^26^ Most fats partition into the n-hexane fraction, while inorganic salts partition into the aqueous part. The advantage of the method is total recovery of the target compounds based on gradual differences in their polarity.^27^ A more recent modification based on the original Kupchan and modified protocol by Wegenen was developed by Muhit, et. al. (2010) for their study for the isolation and identification of different compounds from the leaf extract of *Dillenia indica*. The procedure is illustrated below. For this process methanol serves as the initial solvent. Extraction of the crude methanolic extract with n-hexane yields two layers, an upper n-hexane soluble fraction and a lower aqueous fraction. The aqueous fraction is subjected to further sequential extraction with carbon tetrachloride and chloroform. Each successive extraction should yield an upper aqueous layer that can be subjected to further extraction. The lower carbon tetrachloride and chloroform layers are collected and kept as is.^28^ The advantage of this newer methodology over the older protocols is that it offers a better partitioning process for isolation and identification of various phytochemicals from plant extracts.

A study by Dieu-Hien Truong (2019) showed a significant difference in the extraction yield using different solvents. Among the solvents tested in the study, methanol resulted in the highest extraction yield (33.2%), followed by distilled water (27.0%), ethanol (12.2%), acetone (8.6%), chloroform (7.2%), and dichloromethane (4.9%). This indicates that the extraction efficiency favors the highly polar solvents. Statistical analysis showed that methanol exhibited the properties of an optimal solvent to extract bioactive components, specifically phenolics, flavonoids, alkaloids, and terpenoids. Low levels of flavonoids, alkaloids, and terpenoids were obtained in the water extracts. The study also found that dichloromethane and chloroform had high total terpenoid content. The lowest phenolic, flavonoid, and alkaloid content in the crude extracts were shown by these solvents.^29^

The extraction of various unknown compounds using a liquid-liquid extraction process depends on corresponding polarities of the compounds of concern. It is therefore important to utilize various polar and nonpolar solvents, so as not to exclude other important compounds that may not be extracted with one solvent alone.

### Cytotoxic Activity of *S. saman*

Cytotoxic activity studies on the leaves of *S. saman* are limited. In a review article by Ferdous, et. al., cytotoxic activity of the leaves of *S. saman* has been evaluated as far back as 1982 and 1998. The method used in both of these studies was brine shrimp lethality assay.^22^ It is a method for preliminary screening of cytotoxic potential of a plant extract or other substances, and can be used as a basis for further experiments on animal models.^30^ In the study, hexane, carbon tetrachloride, and dichloromethane soluble fractions of crude methanol extract were tested in vitro against *Artemia salina* for 1 day. Results showed that the cytotoxicity potential exhibited by carbon tetrachloride and dichloromethane soluble fractions were promising when compared with the positive control, vincristine sulfate.^22^ Furthermore, the researchers were able to purify and isolate two compounds from the n-hexane soluble fraction of the methanol extract: lupeol and epilupeol. Another study in the Philippines confirmed the presence of these compounds through nuclear magnetic resonance. These compounds were isolated and identified from the peduncle and leaves, respectively.^31^ Based on a study, lupeol is a natural pentacyclic triterpenoid that acts as an anticancer agent against MCF-7 cells.^32^ It also inhibits LMP1-induced NF-κB activation and reduces NF-κB-dependent LCL viability.^33^

In another study by Azhar, et. al., five kilograms of leaves were extracted with hexane and methanol using a cold extraction method. The extracts were filtered and evaporated to dryness using a rotary evaporator at 40°C. These were then used for brine shrimp bioassay study. The results showed that the methanol extract had better cytotoxic activity than the hexane extract. The LD50 value of *S. saman* leaves of the methanol extract was 757.3672 μg/ml, compared to that of hexane which was >1000 μg/ml.^34^

Shanmugakumars and SatheeshKumar (2014) were able to isolate Pithecolobine-1, the bioactive compound present in the leaves of *S. saman*. The compound was isolated using fractionation of the methanolic extract. This was reported to exhibit good cytotoxic activity within a concentration range of 0.019-0.625 mg/mL.^35^

### Antimitotic Drugs

Antimitotic drugs block cells in mitosis, a type of cell division which results in two daughter cells each with the same number of chromosomes as the parent cell. It has several phases: the interphase (G_1_, S, G_2_) and M phase with each corresponding checkpoints that control and regulate the cell cycle before it can proceed to the subsequent phase. The cycle starts with the G_1_ phase which is a gap period between the synthesis of DNA wherein nutrients and energy are gathered to prepare itself for replication. A cell may stop dividing which is represented by the G_0_ phase. This, however, can be induced to start dividing and undergo the cycle. Before going into the S phase, the cell will evaluate its potential to replicate and also to check for any abnormalities. The most important checkpoint, the restriction checkpoint, is responsible for this process. In the S phase, the cell replicates its DNA to prepare itself to undergo the next phase. The next phase is the G_2_ phase wherein the cell prepares itself for the process of division. Mitosis then occurs in the M phase. Karyokinesis, the division of the nucleus, and cytokinesis, the division of the cell, occur here. The result of this cycle is two daughter cells with chromosomes that are identical to the mother cell.^17^

Vinca alkaloids and taxanes are used in the treatment of cancer. Vinca alkaloids include vincristine, vinblastine, and vinorelbine are agents that inhibit microtubule formation. Vincristine is a chemotherapy drug derived from the Madagascar periwinkle plant in 1959. It is used primarily to treat several types of cancer such as leukemias, lymphomas, neuroblastoma, and sarcomas.^37, 38^ This alkaloid exerts cytotoxic activity by disrupting cellular microtubule formation and arrest cell growth during metaphase. This sequence induces the inhibition of cell replication, including the replication of the cancer cells.^39^

The vinca alkaloids bind to tubulin at the plus-tip of microtubules. In higher concentrations, vincristine induces microtubule depolymerization leading to disruption of mitotic spindle formation; thus cells fail to complete a normal mitosis.^40^

Taxanes on the other hand, bind to a different site on β-tubulin and inhibits its disassembly which result in an arrest in mitosis. The most common taxanes used are docetaxel and paclitaxel. Another drug that acts by inhibiting β-tubulin polymerization is colchicine; it is used in the treatment of gout.^36^

### Sea Urchin

The sea urchin is a widely studied test organism because of its capability to artificially induce spawning, fertilization, and embryogenesis.^41^ Sea urchins were first utilized in pioneering studies evaluating the different effects of carcinogens, among others. In a research by Gutierrez (2016) using *Carica papaya* extract on sea urchin development, papaya extract had shown a significant effect on the mitotic capacity of the embryo. Sea urchin and humans are closely related genetically because they share a similar ancestor, the Deuterostome phyla.^29^

Sea urchins are used extensively in various scientific fields including physiology, embryology, biochemistry, and genetics. Studies involving the use of sea urchins date back as far as around the 19th century, however, these studies are very limited.

In the study of Gutierrez (2016), the specific species of sea urchin used in the experiment was *T. gratilla,* a common sea urchin found in the Philippines. In the study’s methodology, male and female sea urchins were collected and were used to spawn and fertilize the eggs. These were placed in a beaker, and the eggs were then examined for abnormalities. The test used increasing concentration of *C. papaya* extract with colchicine as a positive control. After a few hours, each beaker was examined for the mitotic characteristic of the embryos, and it was shown that the papaya extract had a significant effect on the mitotic capacity of the embryo.^29^

*T. gratilla* is a sea urchin that is classified under the phylum Echinodermata. This species is commonly used in biological testing and assays. It is reported to have bioactive compounds for drug discoveries and pharmacological research. It can also be used as a biological control agent in several procedures.^41–42^

### Operational Definition of Terms

1. Fertilization - starts with the appearance of the fertilization membrane
2. Fertilization membrane - the alteration of the vitelline layer seen as a thickening of the outer border of the embryo
3. Mean cell stage - refers to the average cell stage of 50 cells counted
4. Mitotic inhibition - is based on the average cell stage at the end of the treatment with comparison to the vincristine sulfate control

## CHAPTER III: METHODOLOGY

### Study Design and Time Allocation

In-vitro quasi-experimental study design was used. The research proper ran from August 2019 to February 2020.

### Sampling Area and Collections

Ten live *T. gratilla* were procured from a local fisherman in Barangay Ibo, Lapu-Lapu City, Cebu. Samples were placed in a large cooler box filled with seawater and sand from the site where *T. gratilla* were collected, and transported to Cebu City. An additional six liters of seawater, also from the site of collection, were obtained. The sea urchins were identified using taxonomic keys and was further confirmed with a certification from the Bureau of Fisheries and Aquatic Resources (BFAR) Region 7. Five hundred grams of *S. saman* leaves collected from Compostela, Cebu were speciated at the Coastal Resources and Ecotourism Research Development and Extension Center. Experimental procedure was performed in the Microbiology Laboratory and the Biochemistry Laboratory of the Cebu Institute of Medicine, Cebu City.

### Study Population

#### Inclusion Criteria

Ten live *T. gratilla* individuals were used as test subjects for the experiment. Only fresh *S. saman* leaves were used.

#### Exclusion criteria

The researchers did not include other species of sea urchin and *Samanea*.

#### Sampling procedure

Random sampling method was used in assigning the experimental groups.

### Maneuvers

#### Preparation and Extraction of Leaves

The procedure utilized was adopted from the Modified Kupchan Method for liquid-liquid partition by Muhit, et. al. (2010).^28^

As illustrated in Figure 3, 500 grams of *S. saman* leaves was washed with distilled water and then air-dried. The dried samples were processed in a blender until consistency was powdery. The powdered *S. saman* leaves were soaked in 100% methanol for 48 hours and filtered. The filtrate was concentrated using a rotary evaporator and a crude extract was obtained. 100 ml of methanol solution containing 10 ml of distilled water was then added. The crude methanolic extract was placed into the separatory funnel and 100 ml of the n-hexane was added. The funnel was agitated and allowed to stand undisturbed until complete partition was achieved. The organic portion was collected and the process was repeated three times. The n-hexane fraction was collected and evaporated in vacuo. To the crude methanolic extract that was left, 12.5 ml of distilled water was added and mixed. It was transferred into the separatory funnel and extracted with 100 ml of carbon tetrachloride (CCl_4_). This was repeated three times. This yielded a CCl_4_ fraction and an aqueous solution. The CCl_4_ fraction was collected and evaporated in vacuo. To this solution, 16ml of distilled water was added and mixed. It was then transferred into the separatory funnel and extracted with 100 ml of chloroform (CHCl_3_), repeating the process three times. The CHCl_3_ soluble fraction was collected and evaporated in vacuo. The aqueous solution from the previous CCl_4_ extraction was also evaporated in vacuo. The entire procedure produced four stock solution fractions: hexane, CCl_4_, CHCl_3_, and aqueous.

**Figure 1.**
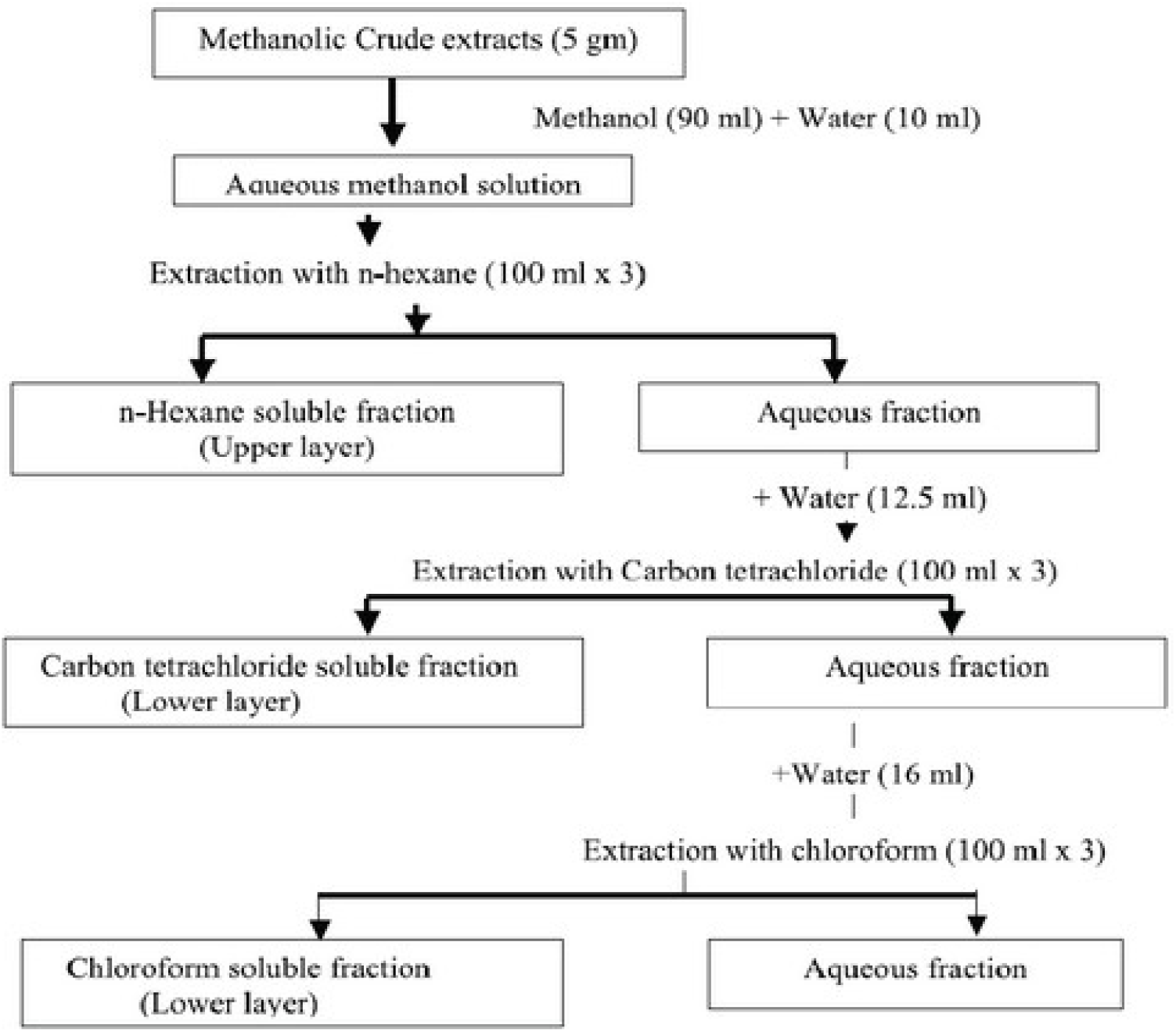
Modified Kupchan’s partition scheme. Adopted from: Muhit et. al., 2010.^28^

**Figure 2.**
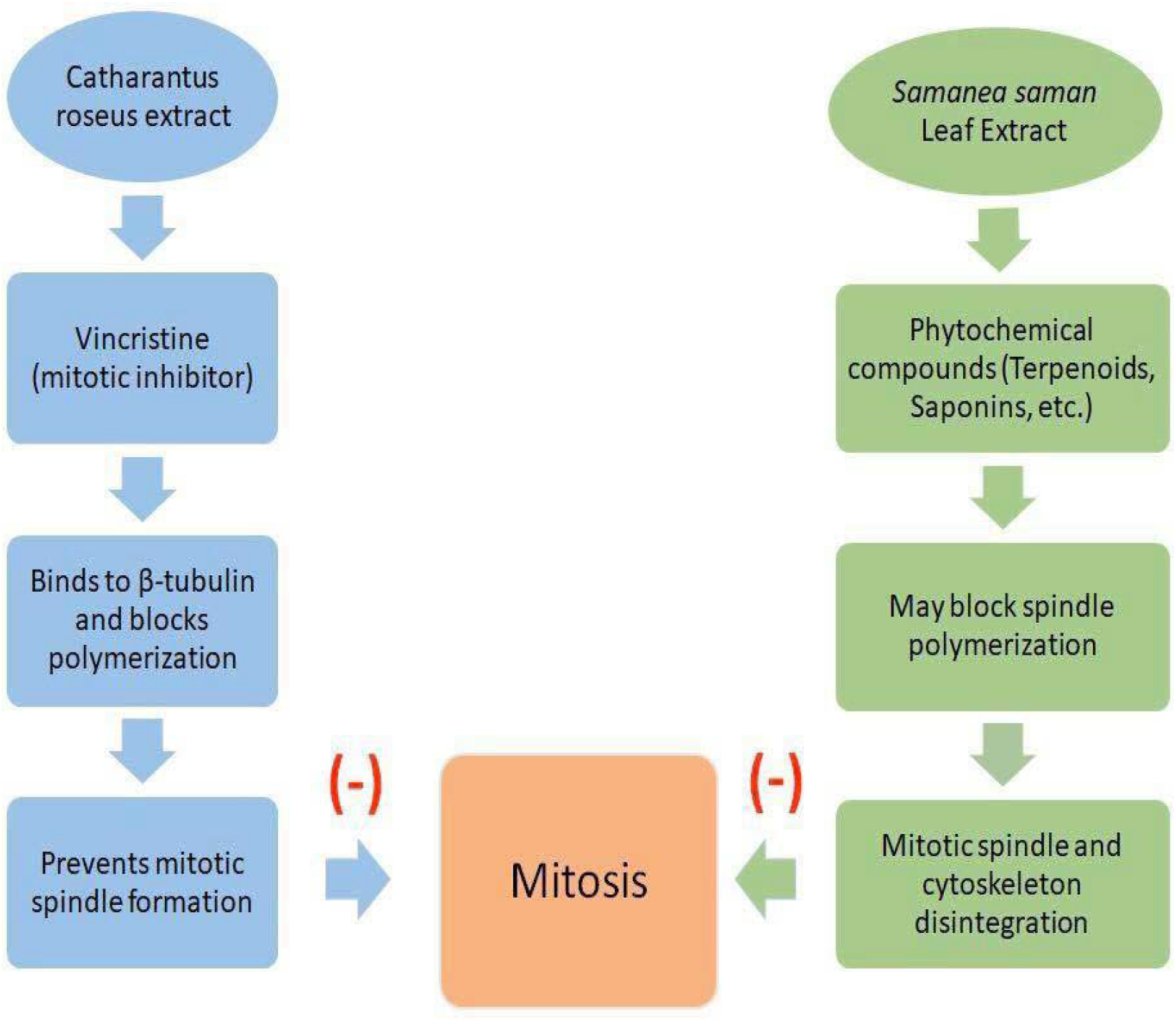
Conceptual framework

**Figure 3.**
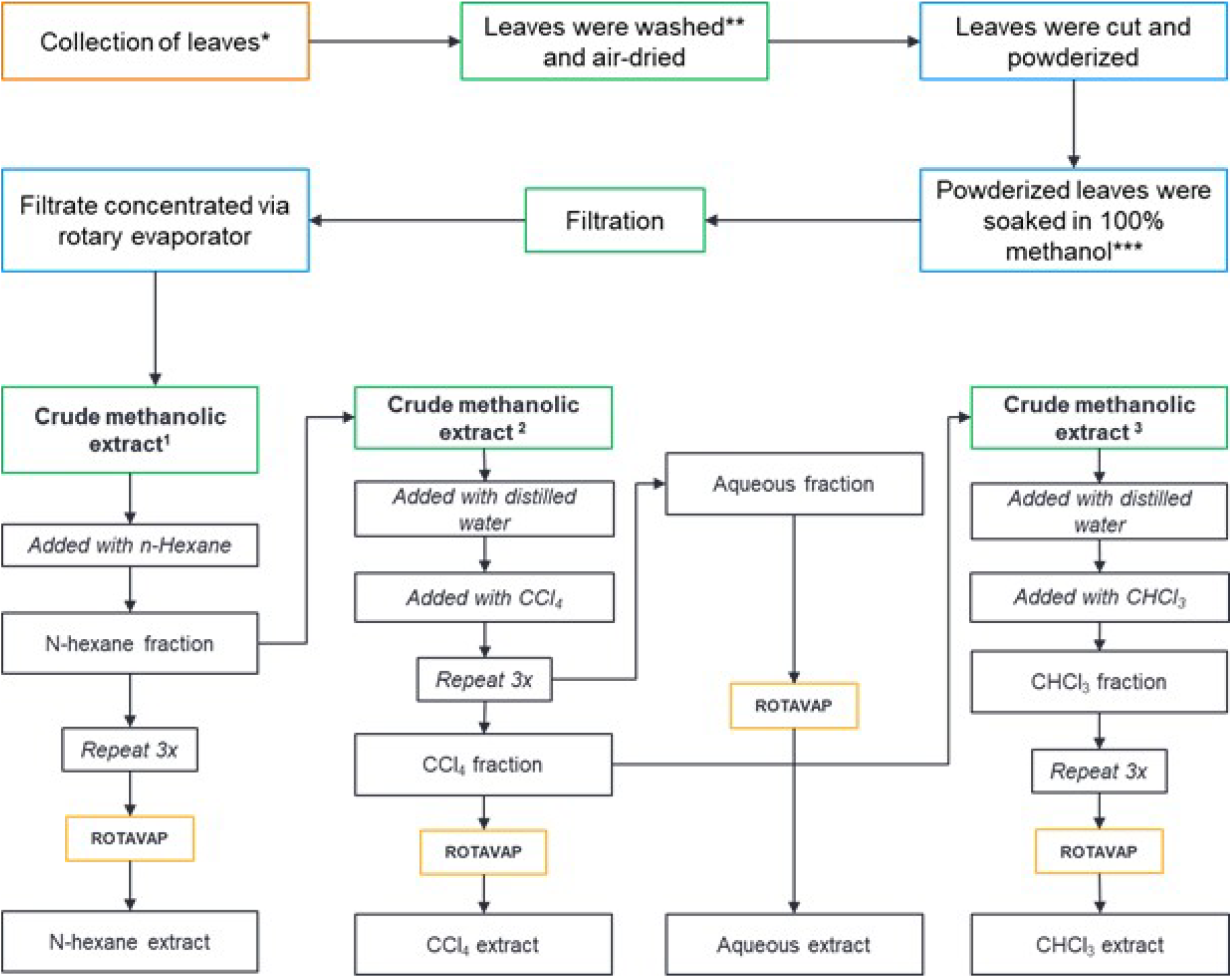
Procedural flowchart for preparation of leaf extracts. Adopted from Muhit, et. al’s Modified Kupchan protocol. *500 grams of leaves; **Distilled water was used; ***Soaked for 48 hours; ^1^Crude methanolic extract after methanolic extraction; ^2^Crude methanolic extract after hexane extraction; ^3^Crude methanolic extract after CCl_4_ extraction

The filtrates were stored in an amber-colored bottle to prevent degradation of any possible photosensitive compounds. The extract was then used to make test solutions by diluting the extract with dimethyl sulfoxide (DMSO).

#### Preparation of Test Solutions

A 1000 ppm stock solution was prepared by dissolving 100 mg of each fraction with 100 ml of DMSO. Different concentrations of the 4 fractions were prepared: 800 ppm, 400 ppm, 200 ppm and 100 ppm. The positive control used was 20 μg/ml vincristine sulfate. The negative control was prepared by adding 4 ml of filtered sea water with 16 ml of DMSO.

#### Sea Urchin Bioassay Ethical Considerations

The ethical review obtained from the IACUC for the use and research of *T. gratilla*. (see Appendix I-A)

#### Collection and Determination of Sex

The *T. gratilla* were placed in seawater-filled beakers. They were segregated by sex which were determined by injecting 1 ml of 0.5 M potassium chloride (KCl) through the peristomial part of the echinoderm. Female *T. gratilla* subjects secreted orange-colored eggs, while male subjects secreted cream-colored sperm cells. The secretions released by the organisms were confirmed microscopically.

As indicated above, in Figure 4, after confirmation of the sea urchins’ sexes, *T. gratilla* sperm cells were collected and placed in beakers. The beakers were placed in an ice water bath at four (4) degrees Celsius to preserve the viability of the sperm sample. A drop of sperm was diluted in 10 ml of filtered seawater and used for fertilization.

**Figure 4.**
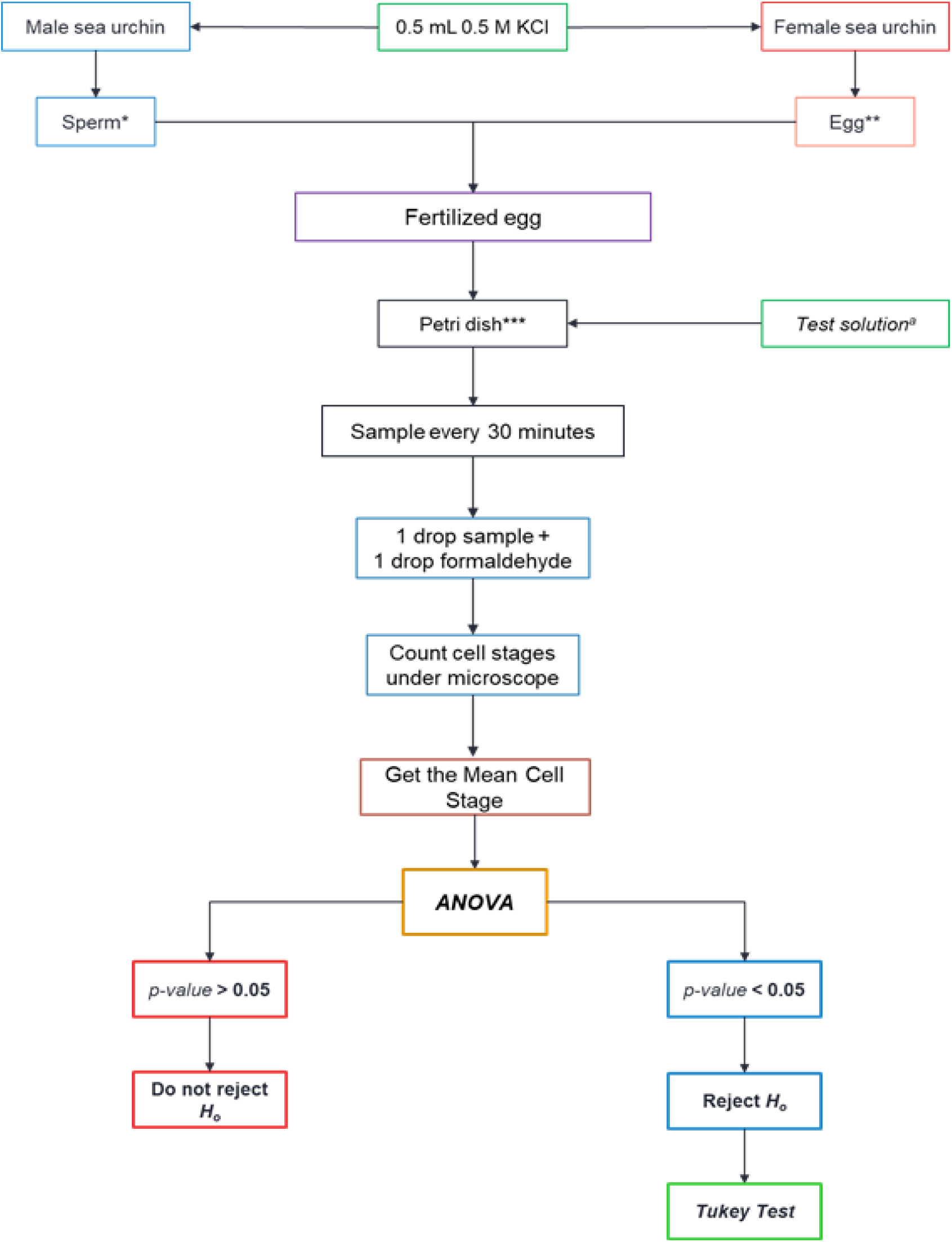
Schematic diagram of the experimental procedure. Adopted from Dajao MA, et al. *in 4°C, 1 drop with 10 mL filtered sea water; **in room temperature, in 500 mL dilution; ***5 mL of fertilized eggs is used in each set-up; ^a^5 mL of each stock solution is used per set-up (*n-hexane*, *CCl_4_*, *CHCl_3_*, and *aqueous* extract). *Sea Urchin Spawning*

Mature eggs of female *T. gratilla* sea urchins, described as cells with small nucleus in the periphery and large cytoplasm, were collected and pooled in a separate beaker filled to the brim with filtered seawater. The eggs were allowed to settle and the excess fluid was drained. This was repeated twice. The eggs were then resuspended in a separate beaker with 500 ml of filtered sea water to finalize the egg suspension.

Fertilization was done by adding 2 ml of diluted sperm to the 500 ml of egg cell suspension. Fertilization was microscopically observed and confirmed by the presence of a fertilization membrane after combining the sperm. Egg suspension of fertilized eggs were then placed in petri dishes.

#### Determination of Antimitotic Activity

Immediately after fertilization, 5 ml of each of the test solutions were added to petri dishes with 5 ml of egg suspension. Each treatment was replicated three times. A drop of sample was removed from each petri dish at 15 minutes after fertilization and every 30 minutes thereafter until the negative control reached a majority of 32-cell stage. 1% formaldehyde was added to the sample to preserve it. A 50-cell count was done using a microscope to check how many cells have gone through the different cell stages. From this the mean cell stage was calculated for each concentration. (see Appendix II-A)

#### Disposal of Used Sea Urchins

The sea urchins were euthanized by freezing for 1 hour and were buried and used as fertilizer afterwards.

### Statistical Analysis

One way-analysis of variance (ANOVA) was used to test significant differences between treatment means. Post hoc test (mean separation test) was conducted using Tukey’s test for the replicates which were significant (p <0.05). Data processing was done using statistical software SPSS-Statistical package. A data collection sheet was used. (see Appendix C)

## CHAPTER 4: RESULTS

The mean cell stage for each treatment was calculated and summarized.

Figure 5 shows that the different hexane and CHCl_3_ fractions are concentration-dependent. Mean cell stages were observed to increase over the different time intervals with the lower concentration treatments of both fractions, specifically Hex 100, CHCl_3_ 800, 400, 200, and 100 ppm. On the other hand, higher concentrations, namely Hex 800, 400, and 200 ppm, were comparable to the positive control with no significant mitotic activity.

**Figure 5.**
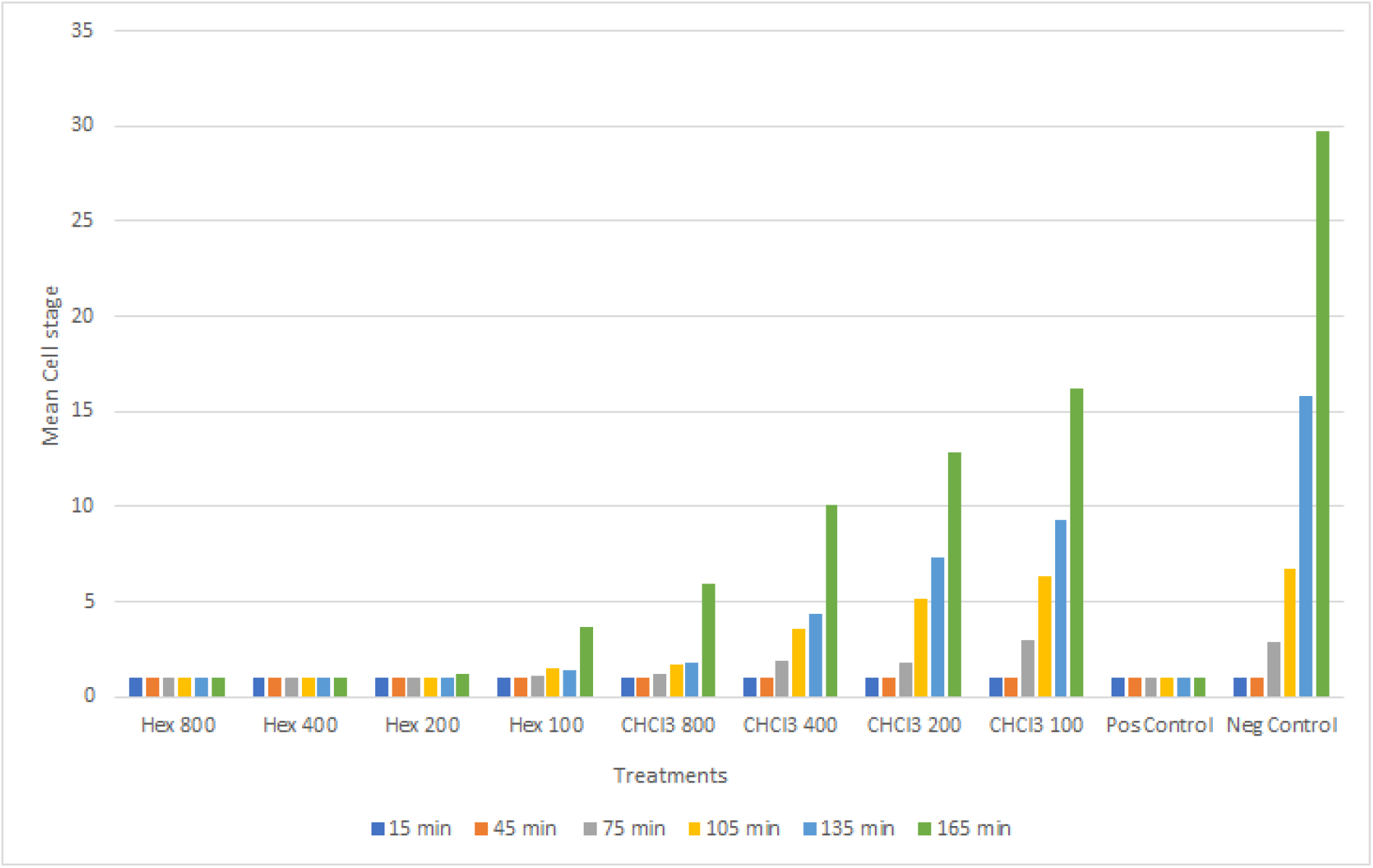
**Graph of hexane and CHCl_3_ treated samples comparing time and mean cell stage**

Similar observations of concentration-dependency of the CCl_4_ and aqueous treatments can be seen in Figure 6. Higher concentration treatments such as the 800, 400, and 200 ppm of both the aqueous and CCl_4_ fractions are comparable to the positive control where little to no mitotic activity was observed. Lower concentration treatments, such as CCl_4_ 100 ppm, showed mitotic activity across the different time intervals.

**Figure 6.**
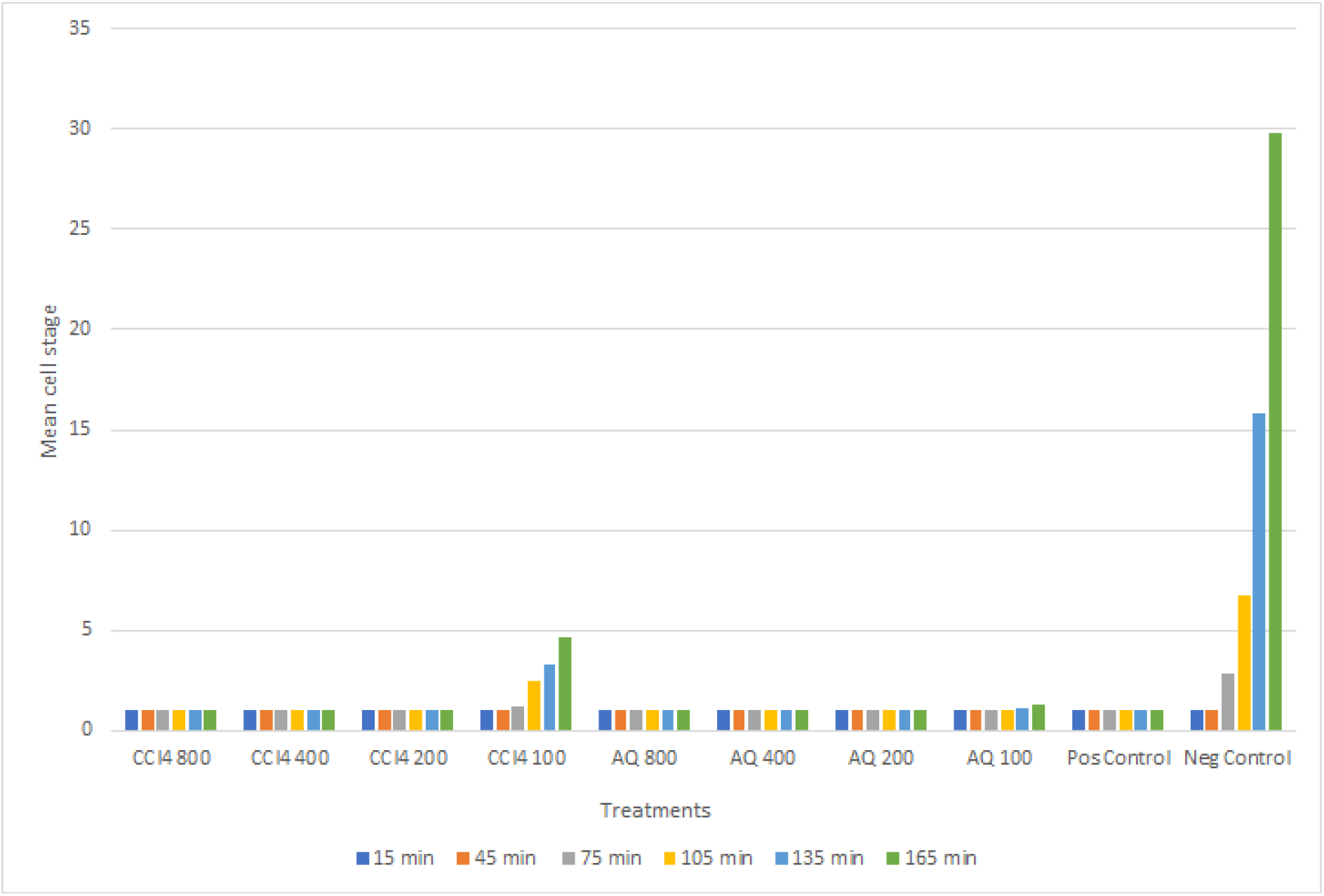
**Graph of CCl_4_ and Aqueous treated samples comparing time and mean cell stage**

One-way ANOVA was used on the mean cell stages of each fraction and their respective concentrations for each time interval (15, 45, 75, 105, 135, and 165 minutes). A Tukey test was done if the null hypothesis was rejected, showing a significant difference between the means.

ANOVA for the treatments at the 15-minute interval was done, and the means showed no significant difference in the mean cell stages across all treatments.

Table 1 shows a p-value greater than 0.05 after the one way ANOVA was done at the 45-minute interval.

**Table 1.** ANOVA at 45 minutes

| **Analysis of Variance (45 min)** |  |  |  |  |  |
| --- | --- | --- | --- | --- | --- |
| **Source** | **DF** | **Adj SS** | **Adj MS** | **F - Value** | **P - Value** |
| **Factor** | 17 | 0.004533 | 0.000267 | * | * |
| **Error** | 36 | 0 | 0 |  |  |
| **Total** | 53 | 0.00453 |  |  |  |

At the 75-minute interval, a p value of <0.05 was obtained after ANOVA (Table 2) showing a significant difference between the means of the different extracts. A post-hoc Tukey test (see Appendix D-IIA) was then done accordingly.

**Table 2.**
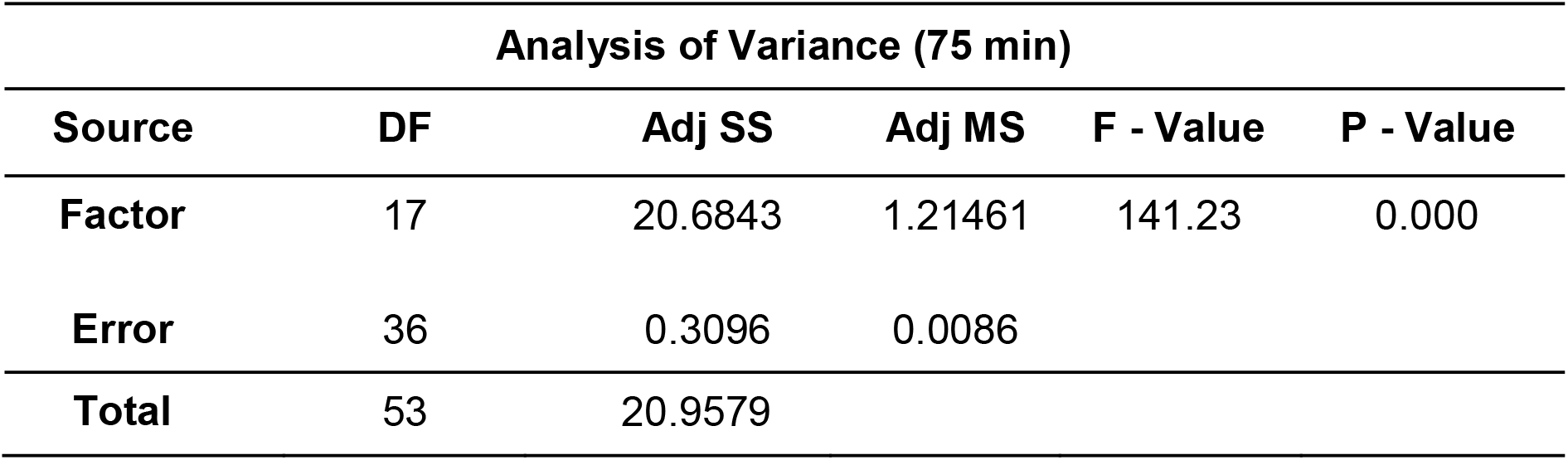
ANOVA at 75 minutes

| Analysis of Variance (75 min) |  |  |  |  |  |
| --- | --- | --- | --- | --- | --- |
| Source | DF | Adj SS | Adj MS | F - Value | P - Value |
| Factor | 17 | 20.6843 | 1.21461 | 141.23 | 0.000 |
| Error | 36 | 0.3096 | 0.0086 |  |  |
| Total | 53 | 20.9579 |  |  |  |

CHCl_3_ 100, 200, and 400 ppm fractions showed a significant difference compared to the positive control (Table 3). There was no significant difference for hexane and CCl_4_ fractions of all concentrations, and aqueous fractions of 800, 400, and 200 ppm concentrations.

**Table 3.** Tukey test at 75 minutes compared to the positive control (vincristine sulfate)

| No significant difference | Significant difference |
| --- | --- |
| Hexane<br>100, 200, 400, 800 ppm<br>CCl<sub>4</sub><br>100, 200, 400, 800 ppm<br>Aqueous<br>200, 400, 800 ppm | CHCl<sub>3</sub><br>100, 200, 400 ppm |

A p value of <0.05 was obtained at the 105-minute interval (Table 4) following ANOVA indicating a significant difference between all the means. A Tukey test was then done accordingly (see Appendix D-IIB).

**Table 4.** ANOVA at 105 minutes

| **Analysis of Variance (105 min)** |  |  |  |  |  |
| --- | --- | --- | --- | --- | --- |
| **Source** | **DF** | **Adj SS** | **Adj MS** | **F - Value** | **P - Value** |
| **Factor** | 17 | 194.611 | 11.4477 | 632.08 | 0.000 |
| **Error** | 36 | 0.652 | 0.0181 |  |  |
| **Total** | 53 | 195.263 |  |  |  |

At 105 minutes, there was a significant difference between the positive control and all concentrations of the CHCl_3_ fractions as well as the 100 ppm concentrations of both the hexane and CCl_4_ fractions (Table 5). In contrast, all concentrations of the aqueous fraction, and the hexane and CCl4 fractions of 200, 400, and 800 ppm concentrations showed no significant difference.

**Table 5.** Tukey test at 105 minutes compared to the positive control (vincristine sulfate)

| **No significant difference** | **Significant difference** |
| --- | --- |
| Hexane | Hexane |
| 200, 400, 800 ppm | 100 ppm |
| CCl<sub>4</sub> | CCl<sub>4</sub> |
| 200, 400, 800 ppm | 100 ppm |
| Aqueous | CHCl<sub>3</sub> |
| 100, 200, 400, 800 ppm | 100, 200, 400, 800 ppm |

Table 6 shows that a p value of <0.05, indicating a significant difference between the means off the extracts, was obtained at the 135-minute mark. As with the previous time interval, a Tukey test was performed (see Appendix D-IIC).

**Table 6.** ANOVA at 135 minutes

| Analysis of Variance (135 min) |  |  |  |  |  |
| --- | --- | --- | --- | --- | --- |
| Source | DF | Adj SS | Adj MS | F - Value | P - Value |
| Factor | 17 | 818.618 | 48.154 | 513.37 | 0.000 |
| Error | 36 | 30377 | 0.0938 |  |  |
| Total | 53 | 821.995 |  |  |  |

The results of the Tukey test (Table 7) show that there was a significant difference between the positive control and the 100, 200, and 400 ppm concentrations of the CHCl_3_ fraction as well as the CCl_4_ fraction 100 ppm concentration. On the other hand all concentrations of the hexane and aqueous fractions, the 200, 400, and 800 ppm concentrations of the CCl4 fraction, and the 800 ppm concentration of the CHCl_3_ fraction showed no significant difference.

**Table 7.** Tukey Test at 135 minutes compared to the positive control (vincristine sulfate)

| No significant difference | Significant difference |
| --- | --- |
| CHCl<sub>3</sub> | CHCl<sub>3</sub> |
| 800 ppm | 100, 200, 400 |
| CCl<sub>4</sub> | CCl<sub>4</sub> |
| 200, 400, 800 ppm | 100 |
| Hexane |  |
| 100, 200, 400, 800 ppm |  |
| Aqueous |  |
| 100, 200, 400, 800 ppm |  |

Table 8 shows that a p value of <0.05 was obtained at the 165-minute interval. As this indicates a significant difference between all the means thus rejecting the null hypothesis, a Tukey test was performed (see Appendix D-IID).

**Table 8.** ANOVA at 165 minutes

| Analysis of Variance (165 min) |  |  |  |  |  |
| --- | --- | --- | --- | --- | --- |
| Source | DF | Adj SS | Adj SS2 | F - Value | P - Value |
| **Factor** | 17 | 2922.77 | 171.927 | 522.65 | 0.000 |
| **Error** | 36 | 11.84 | 0.329 |  |  |
| **Total** | 53 | 2934.61 |  |  |  |

At the 165-minute interval, the 100 ppm concentrations of both the hexane and CCl_4_ fractions as well as all concentrations of the CHCl_3_ fraction showed significant difference compared to the positive control (Table 9).

**Table 9.** Tukey Test at 165 minutes compared to the positive control (vincristine sulfate)

| No significant difference | Significant difference |
| --- | --- |
| Hexane<br>200, 400, 800 ppm | Hexane<br>100 ppm |
| CCl<sub>4</sub><br>200, 400, 800 ppm | CCl<sub>4</sub><br>100 ppm |
| Aqueous<br>100, 200, 400, 800 ppm | CHCl<sub>3</sub><br>100, 200, 400, 800 ppm |

## DISCUSSION

As shown in Figure 5 and 6, the treatments were observed to be concentration-dependent—showing higher restriction in mitotic activity at higher concentrations. Treatments that were of 100 ppm in concentration, except that of the aqueous fraction, consistently showed a significant difference to that of the positive control. This indicates that lower concentrations were not as effective in arresting mitosis with respect to that of the vincristine sulfate. In contrast, treatments of higher concentrations, except for the CHCl_3_ fractions, consistently showed no significant difference to the positive control based on Tukey test. Aqueous and CCl_4_ fractions demonstrated considerable antimitotic properties the most, showing inhibition across all concentrations.

*S. saman* leaf extracts are found to contain terpenoids and saponins reported to have cytotoxic activity in some studies.^14, 31–33^ These compounds are dependent on the polarity of the extracting solvent. In this study, polar compounds, such as saponins, were more likely to be in the aqueous fractions, whereas nonpolar compounds, such as terpenoids, were more likely to be in the hexane fractions.^42–43^ Compounds that are in between in polarity may have been found in across all four fractions or in a combination of the different fractions. This may have been responsible for the difference in antimitotic activity for each fraction.

Mitosis is inhibited by vincristine sulfate by binding to microtubules thus, preventing polymerization and subsequent cell division. The treatments were based on phytochemical studies that showed that crude leaf extracts of the *S. saman* leaves exhibit tannins, flavonoids, steroids, saponins, cardiac glycosides, and terpenoids which may have antimitotic capabilities. Thus, the extracts may have had a similar mechanism of action to that of vincristine sulfate in the inhibition of mitosis of the *T. gratilla* embryo.^14^ Other mechanisms that may have taken place are by inhibition of cAMP formation, and direct toxicity to the cells among others.^45^

The aqueous and CCl_4_ extracts showed inhibition at 100 ppm, the lowest concentration in this study. Inhibition may possibly still be demonstrated at even lower concentrations, however, due to technical limitations the concentration was restricted only up to 100 ppm. More studies on these extracts with a wider range of concentration would possibly yield better results.

The foremost limitation of this study is that phytochemical studies were out of scope for this paper. The presence of certain phytochemical constituents reported to have cytotoxic activity in *S. saman* leaves could not be confirmed.

Financial and technical limitations restricted this study to an in vitro testing on sea urchin models. Moreover, the pharmacokinetics and dynamics of the metabolites extracted are unknown due to the aforementioned in vitro nature. Factors affecting the sea urchins’ environment such as pH, salinity, and other seasonal variations were also not accounted for.

## CONCLUSION

The extracts of *S. saman* (aqueous 100, 200, 400, 800 ppm and hexane 200, 400, 800 ppm and CCl_4_ 200, 400, 800 ppm) possess potential antimitotic activity against *T. gratilla* embryos comparable to vincristine sulfate as positive control. Thus, it is a potential alternative to vincristine sulfate as an antimitotic agent.

## RECOMMENDATION

The researchers recommend that further studies be conducted for proper identification and isolation of the specific biologic compound responsible for mitotic inhibition. The specific compound should be extracted and retested for its ability. Testing on vertebrate embryos and cells, and possibly human cancer cell lines would be the next step to determine its efficacy as a potential alternative drug. The researchers also recommend finding the right amount of dose that would not be toxic on normal cells but would be efficacious on abnormal or cancerous cells.

## APPENDIX A: Certificates and Letters

### I. Certificate of Approval from IACUC

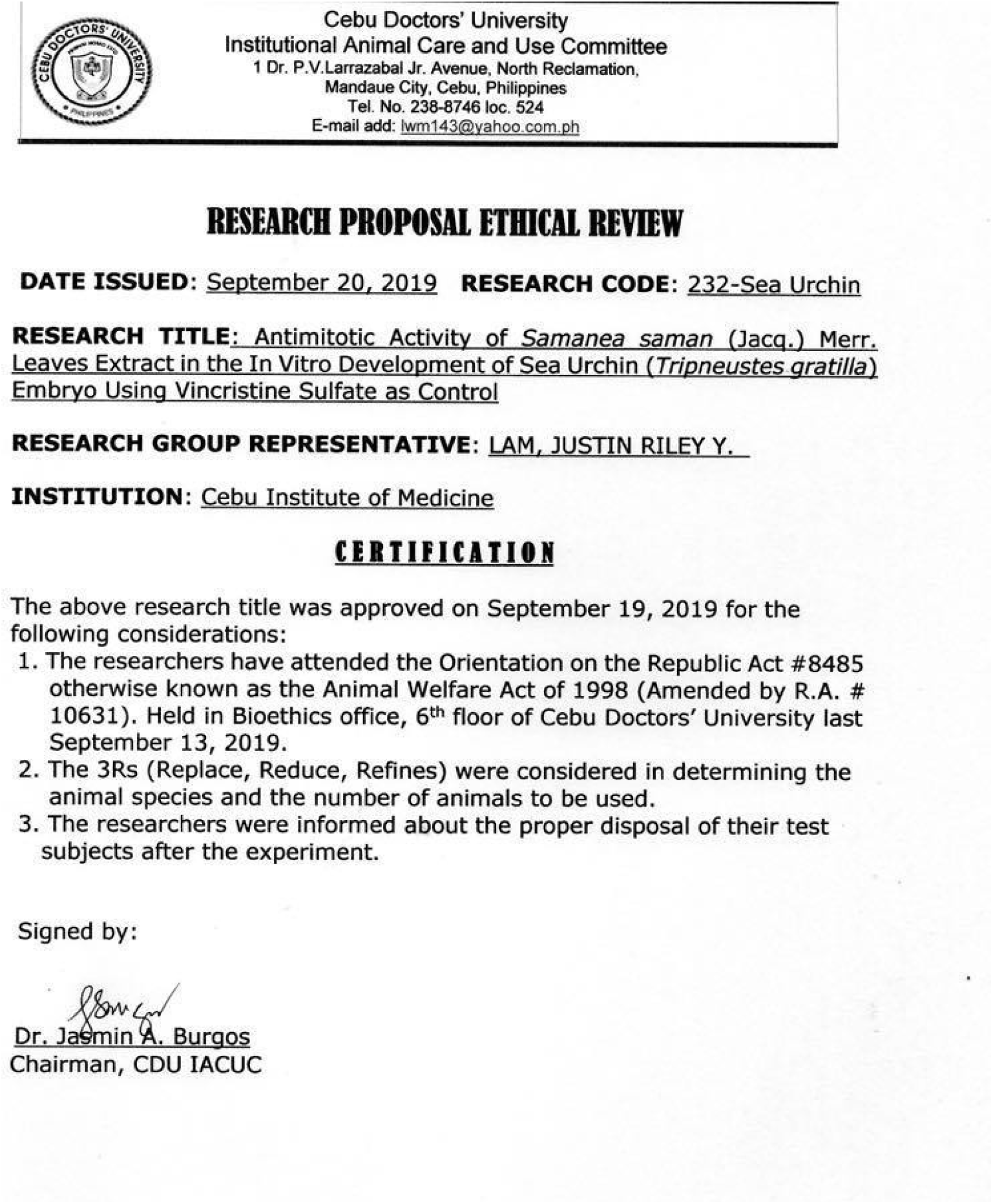

### II. Certification of *Samanea saman*

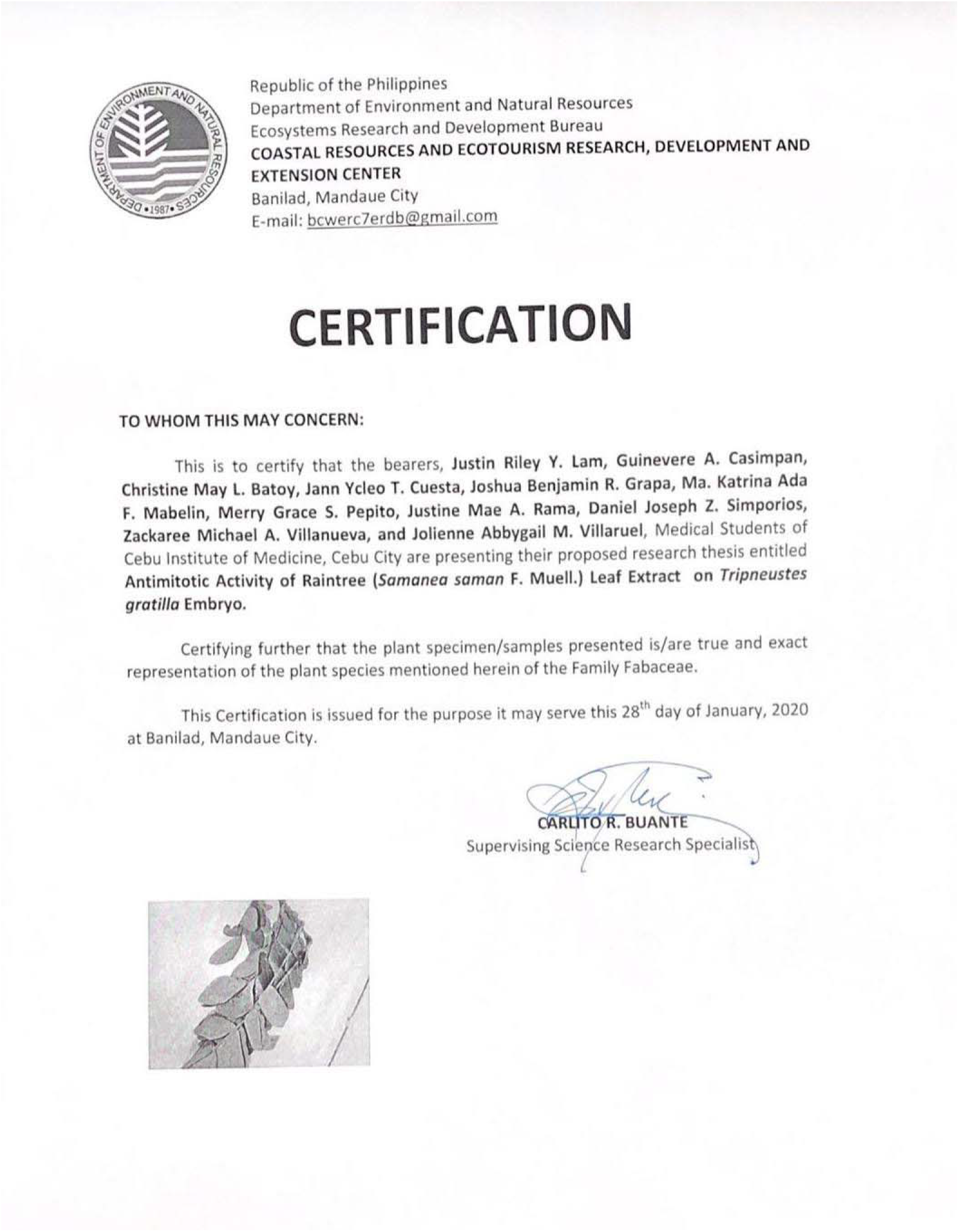

### III. Transmittal Letter for Application for Authorization and Protocol Review for IACUC

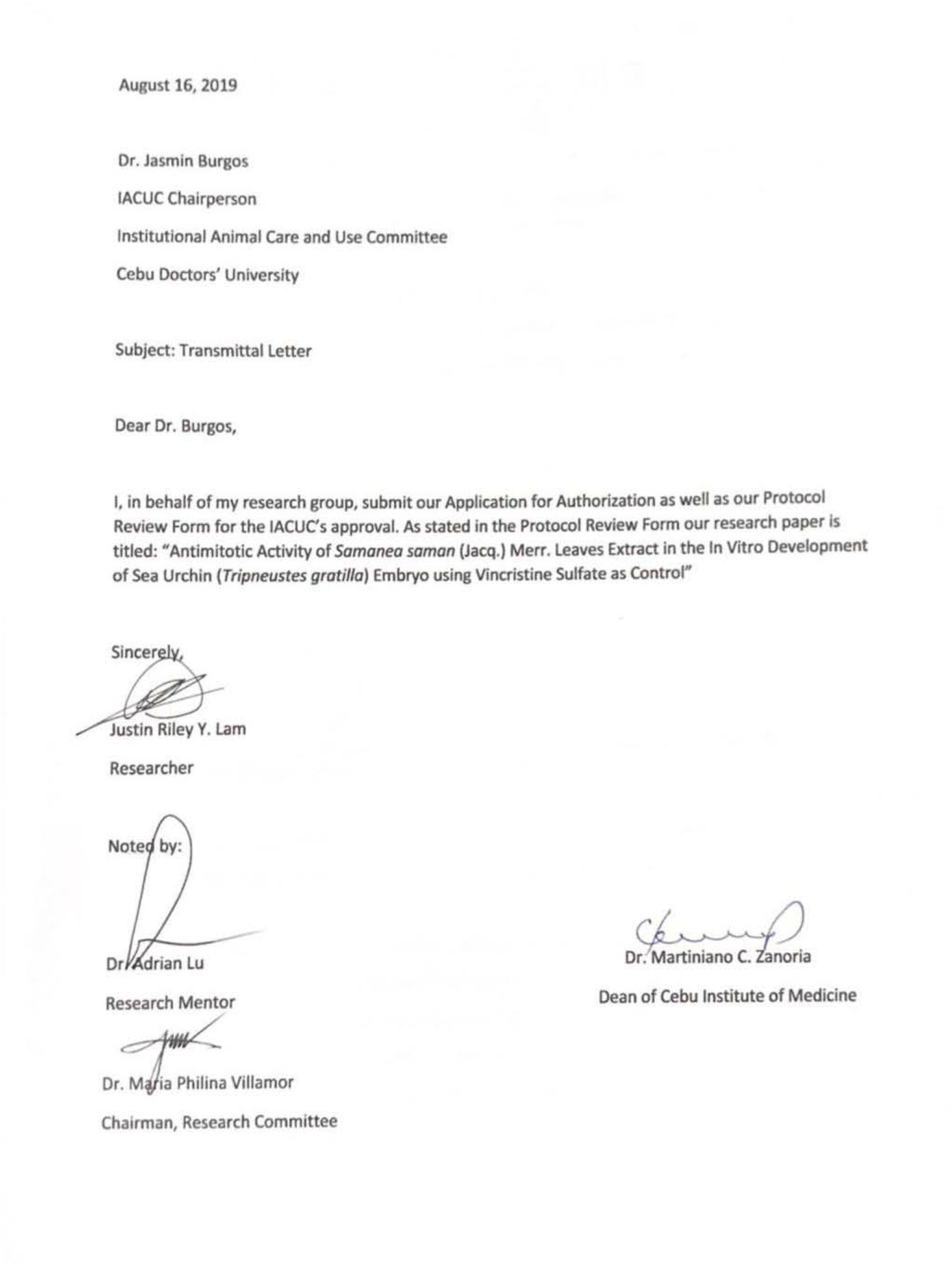

### IV. Application for Authorization and Protocol Review for IACUC

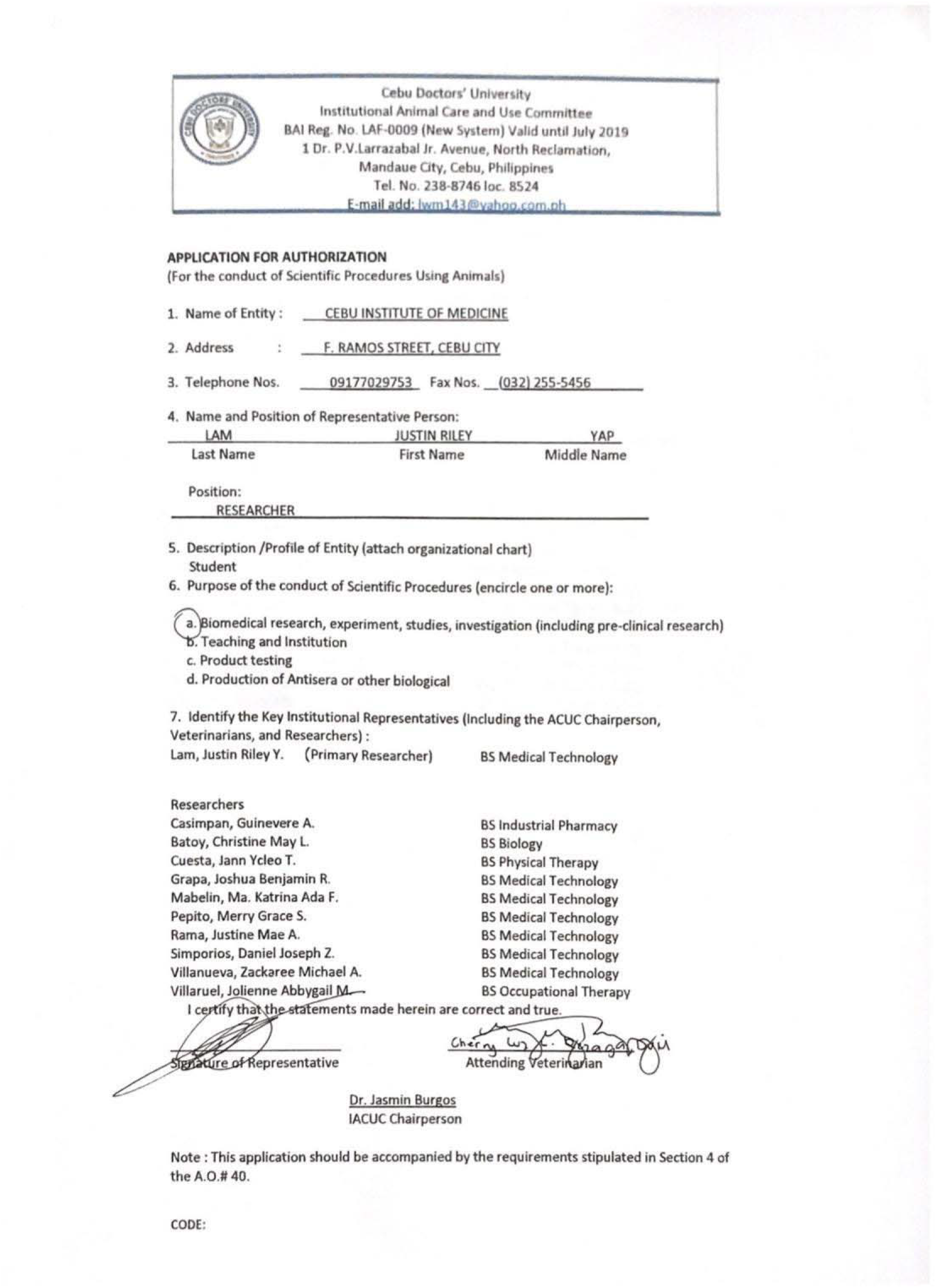
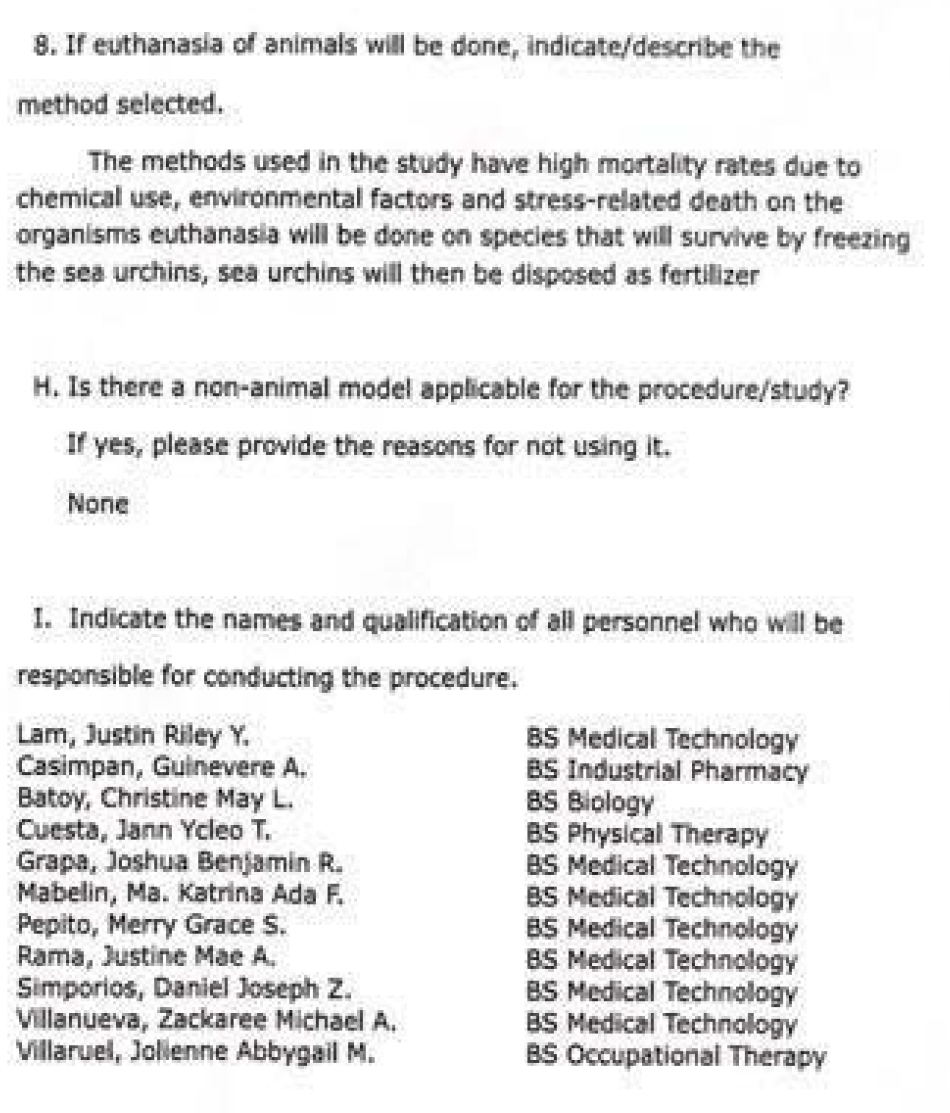

### V. Letter for Use of Laboratory and Materials

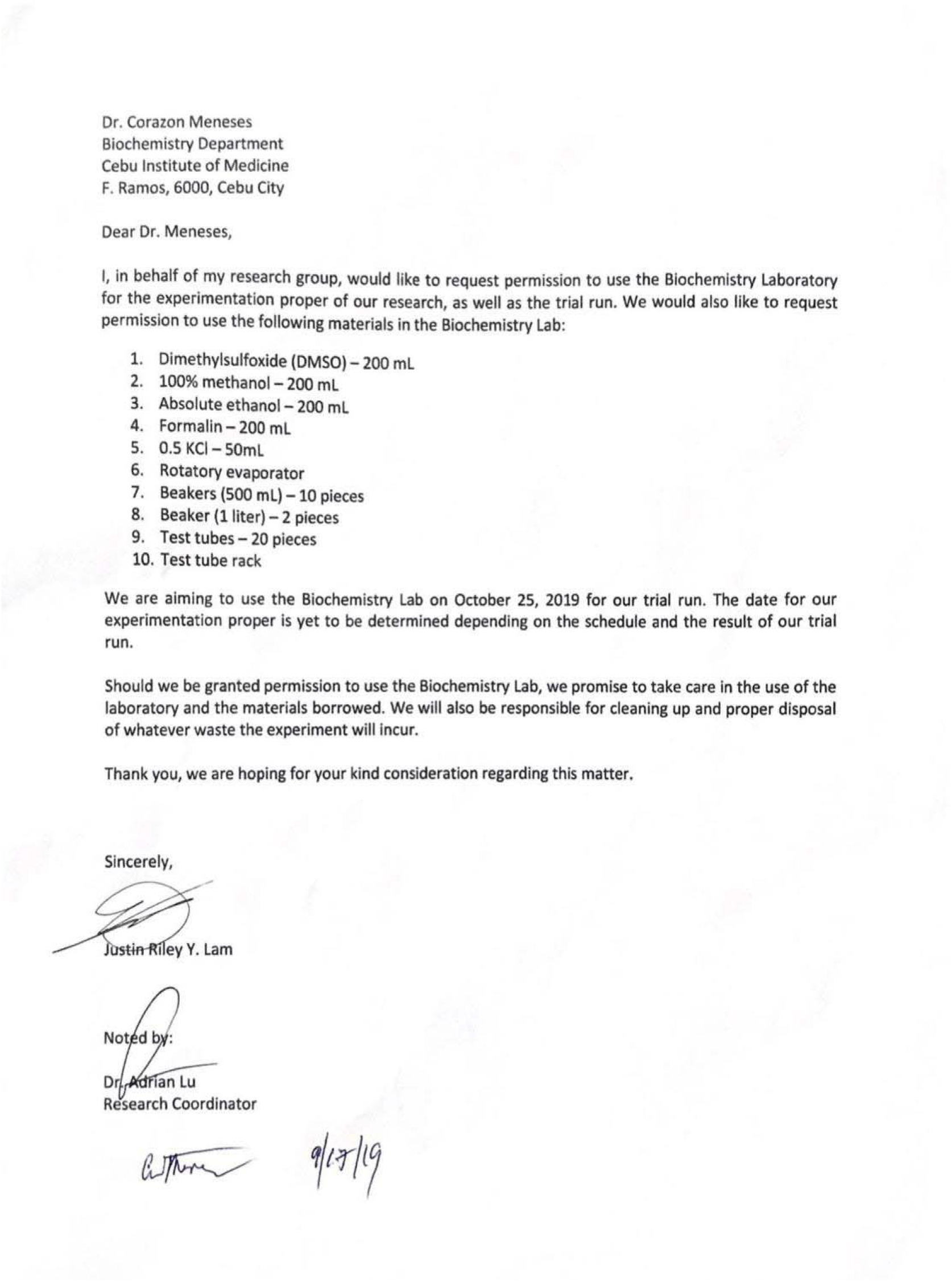

### VI. Letter for Procurement of Hexane

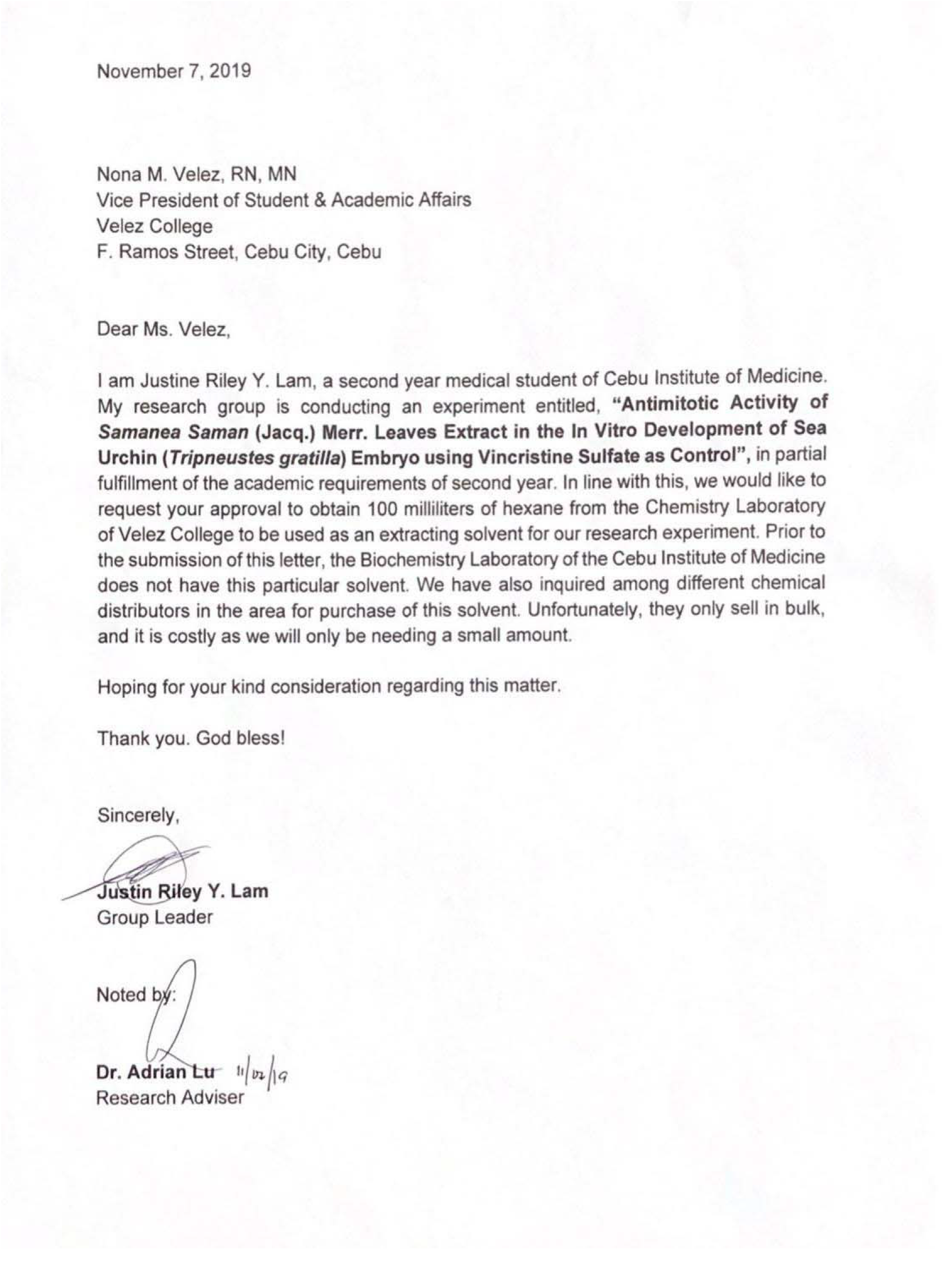

### VII. Letter for Use of Rotary Evaporator and Phytochemical Testing

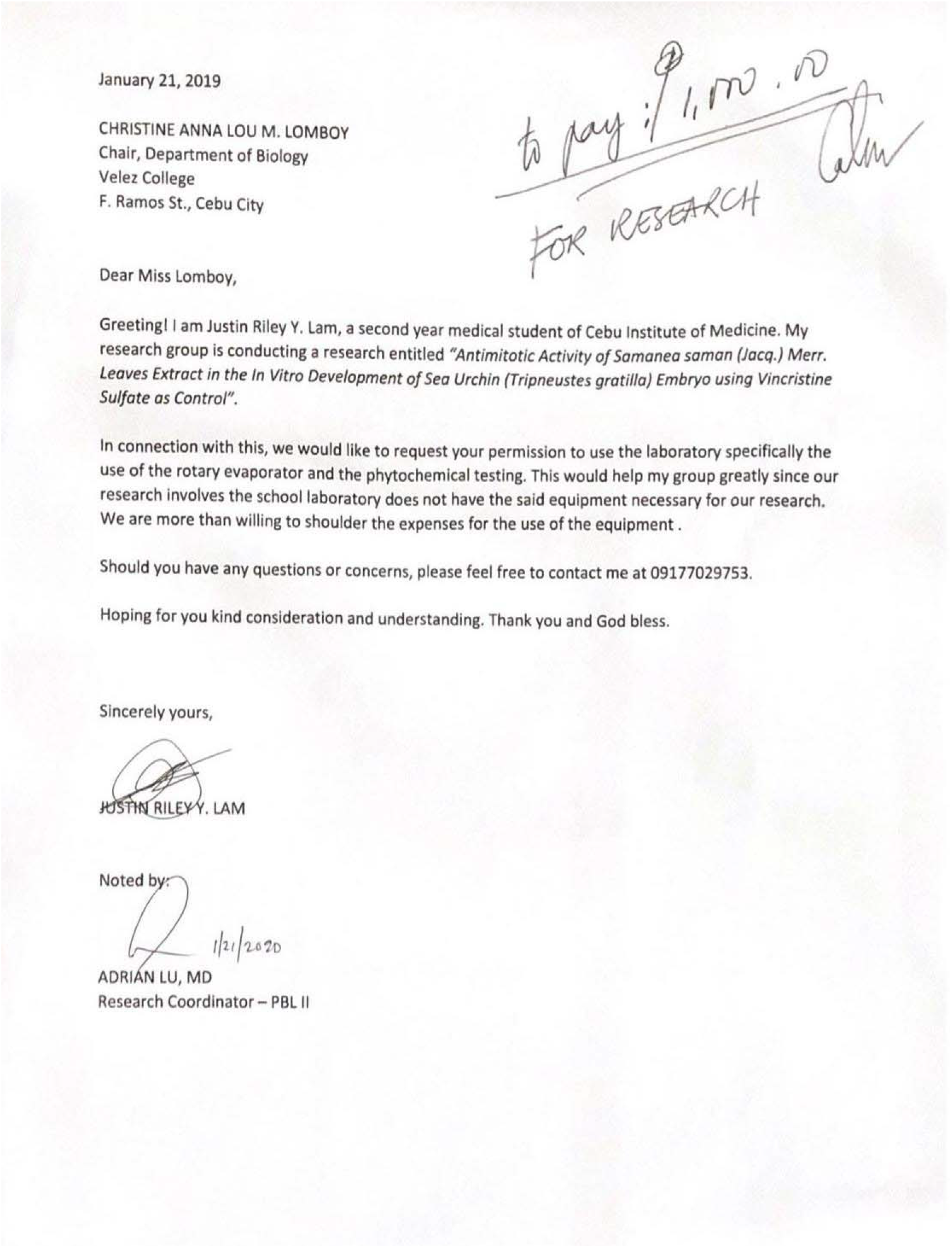

### VIII. Letter for Procurement of Chloroform

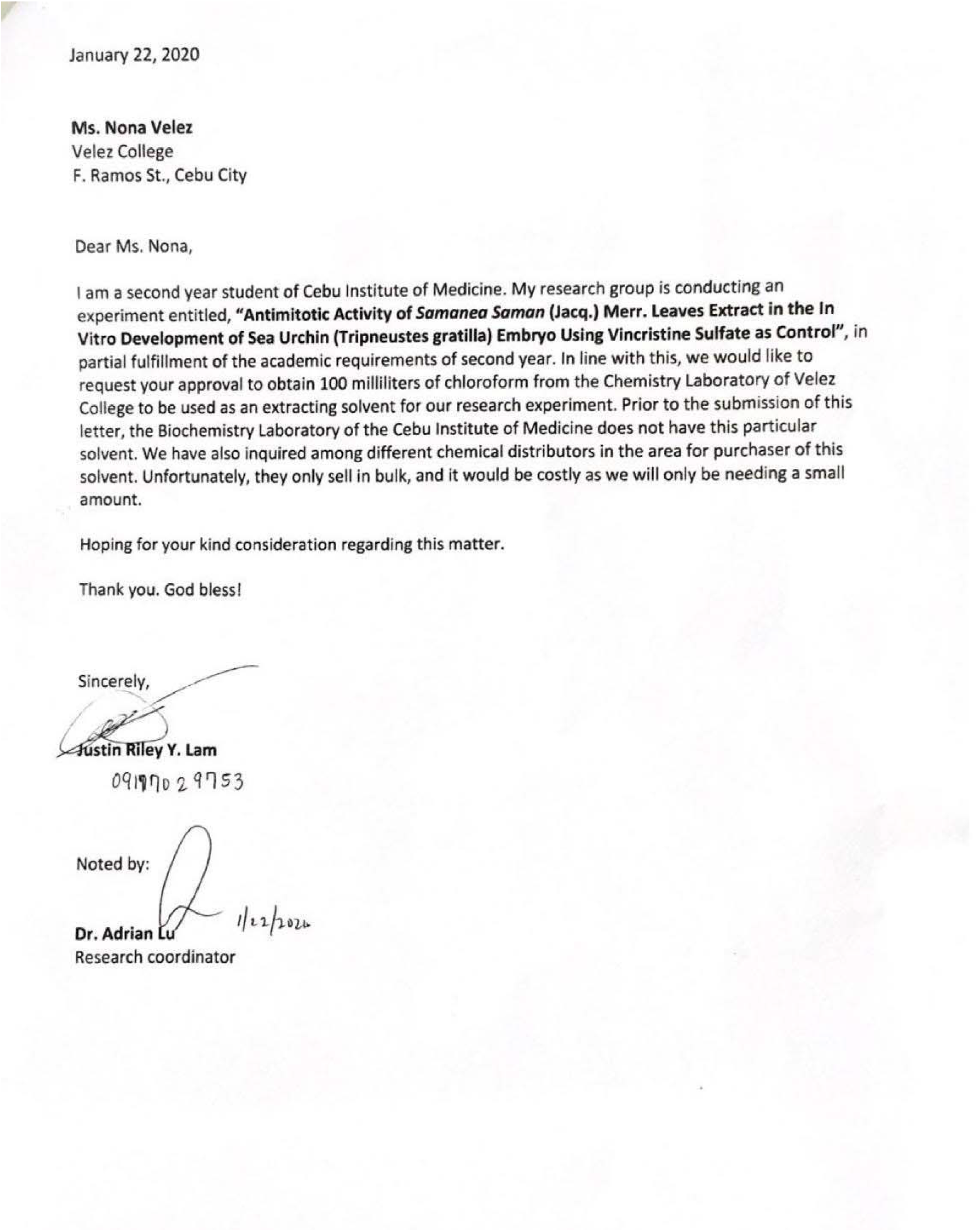

## APPENDIX B: Computation of the Mean Cell Stage

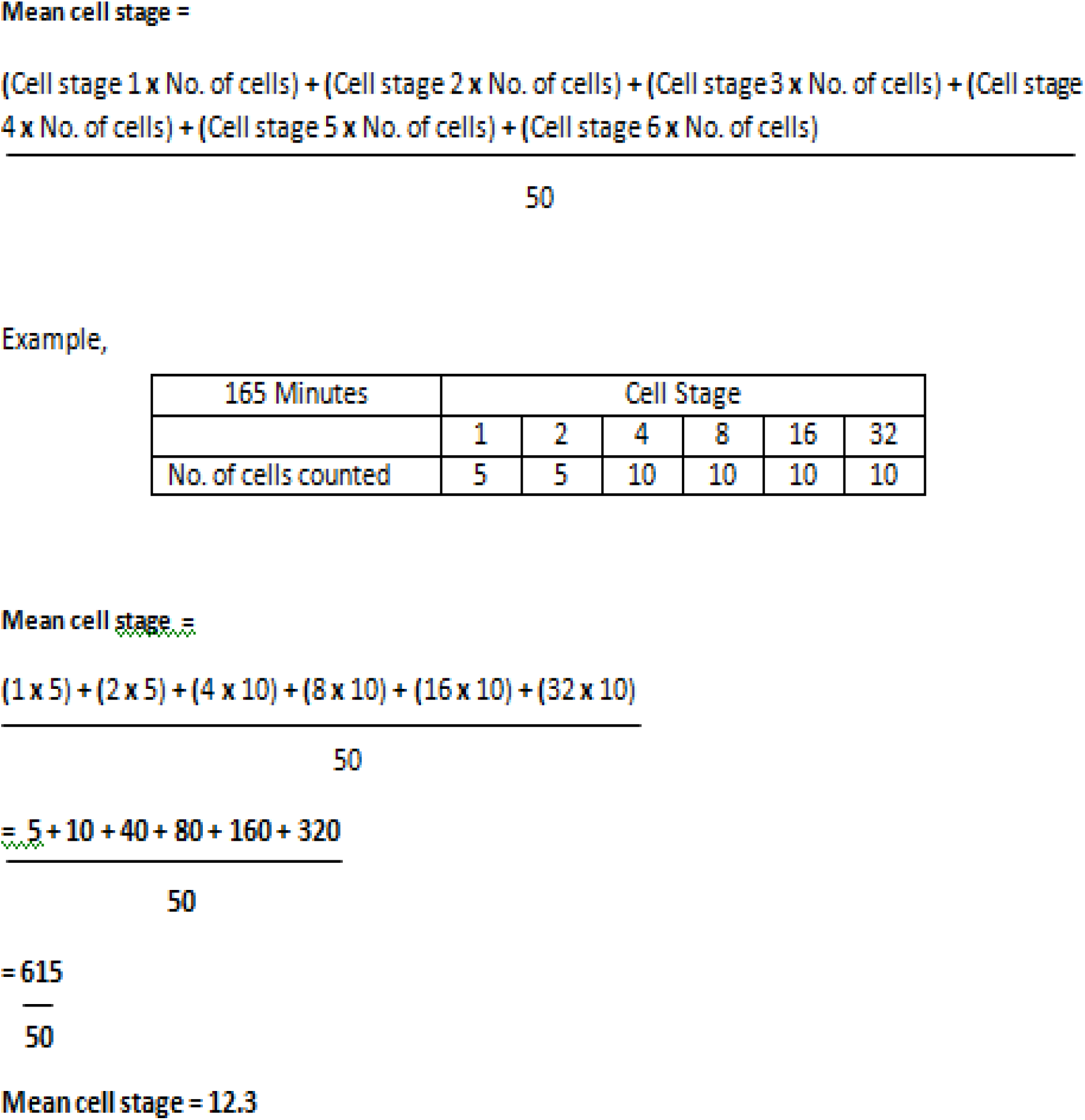

## APPENDIX C: Raw Data

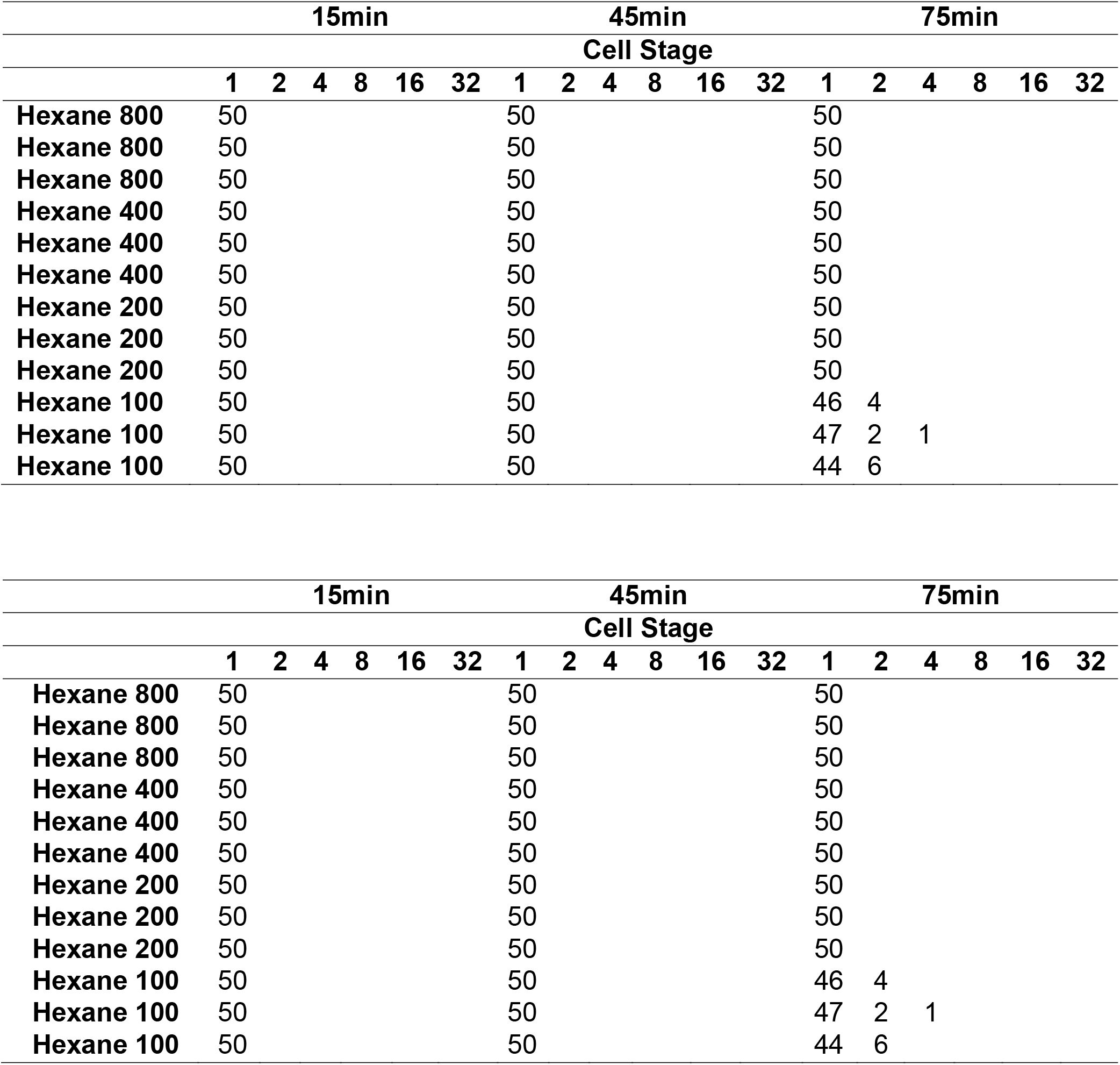
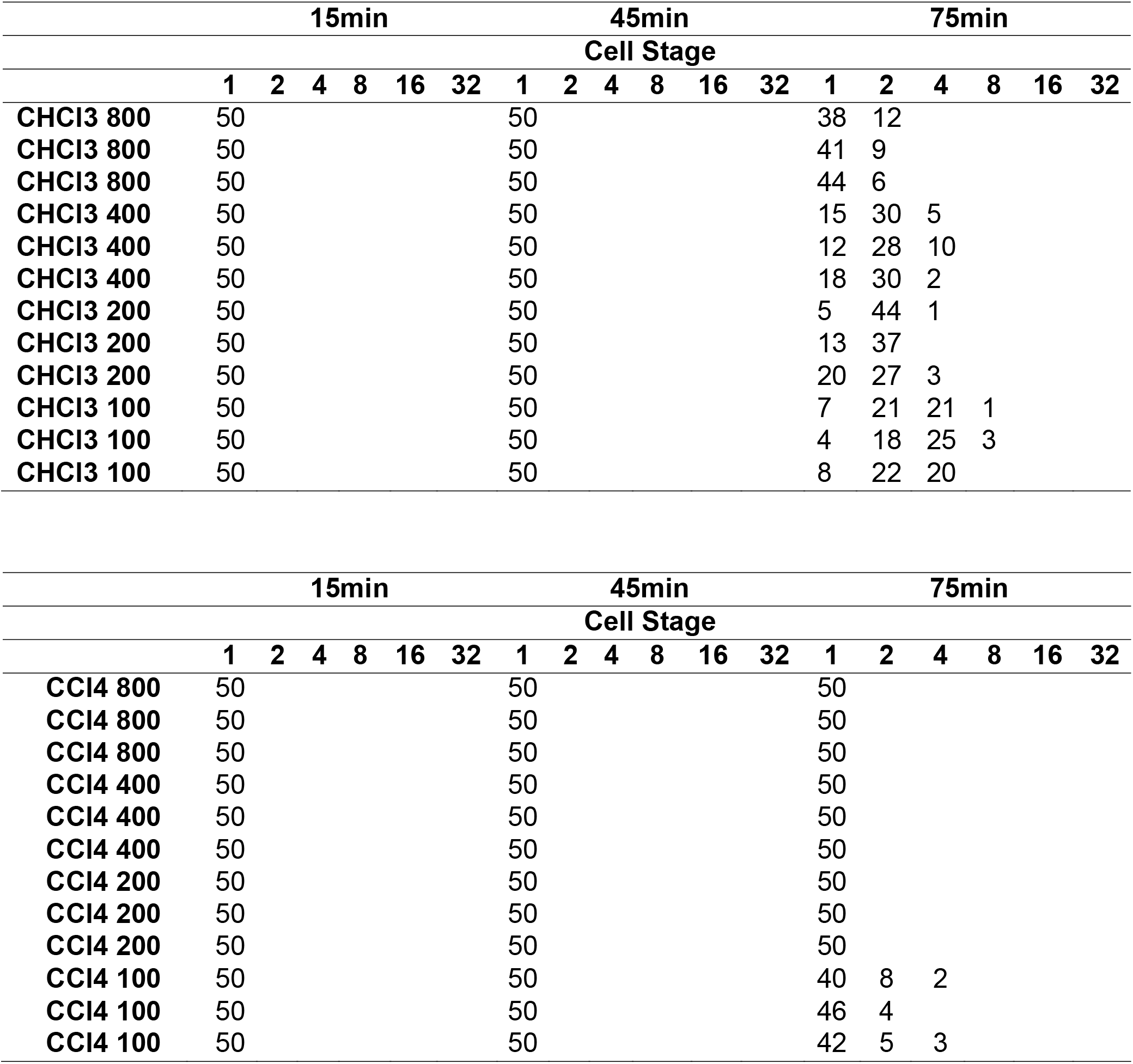
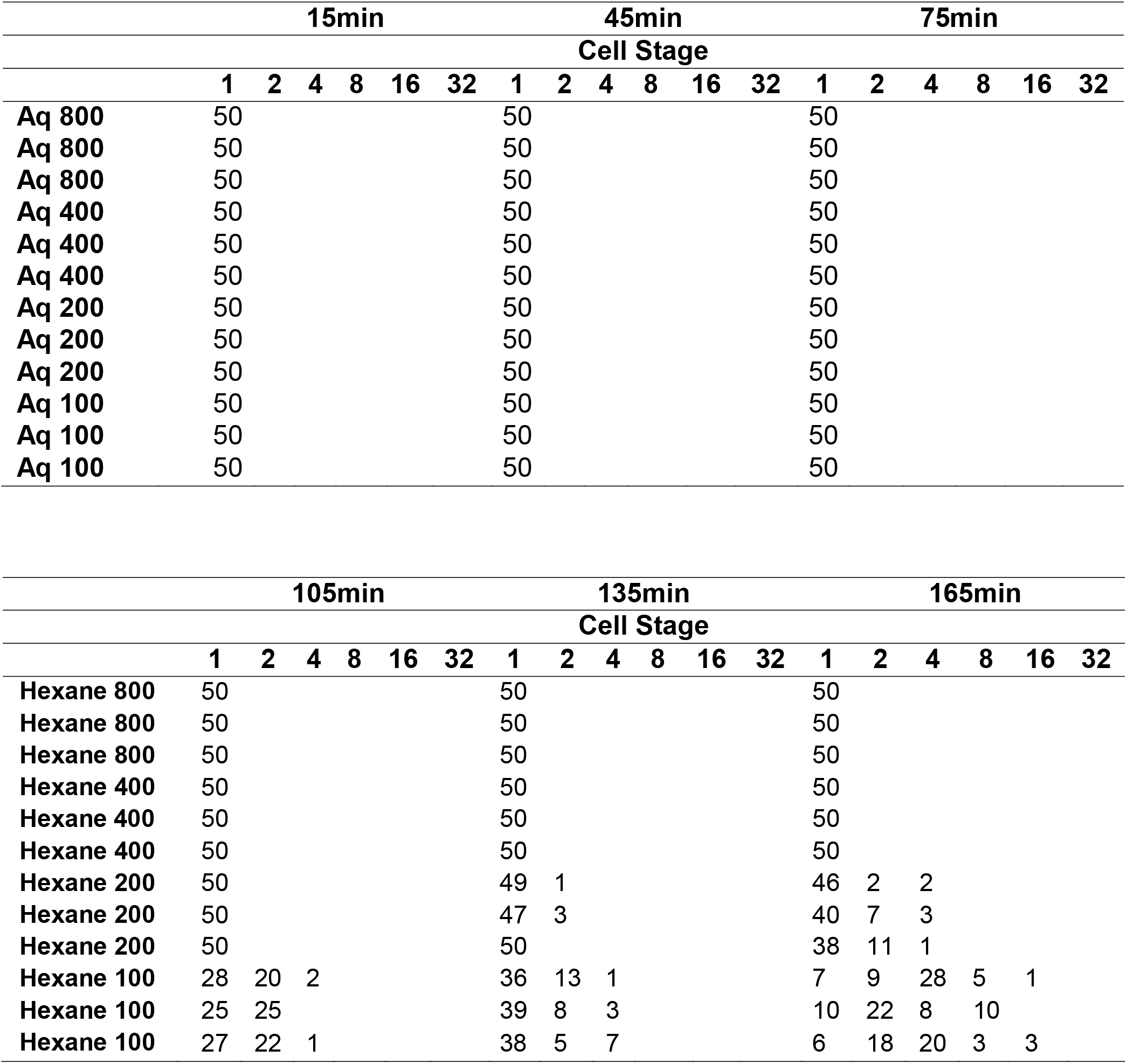
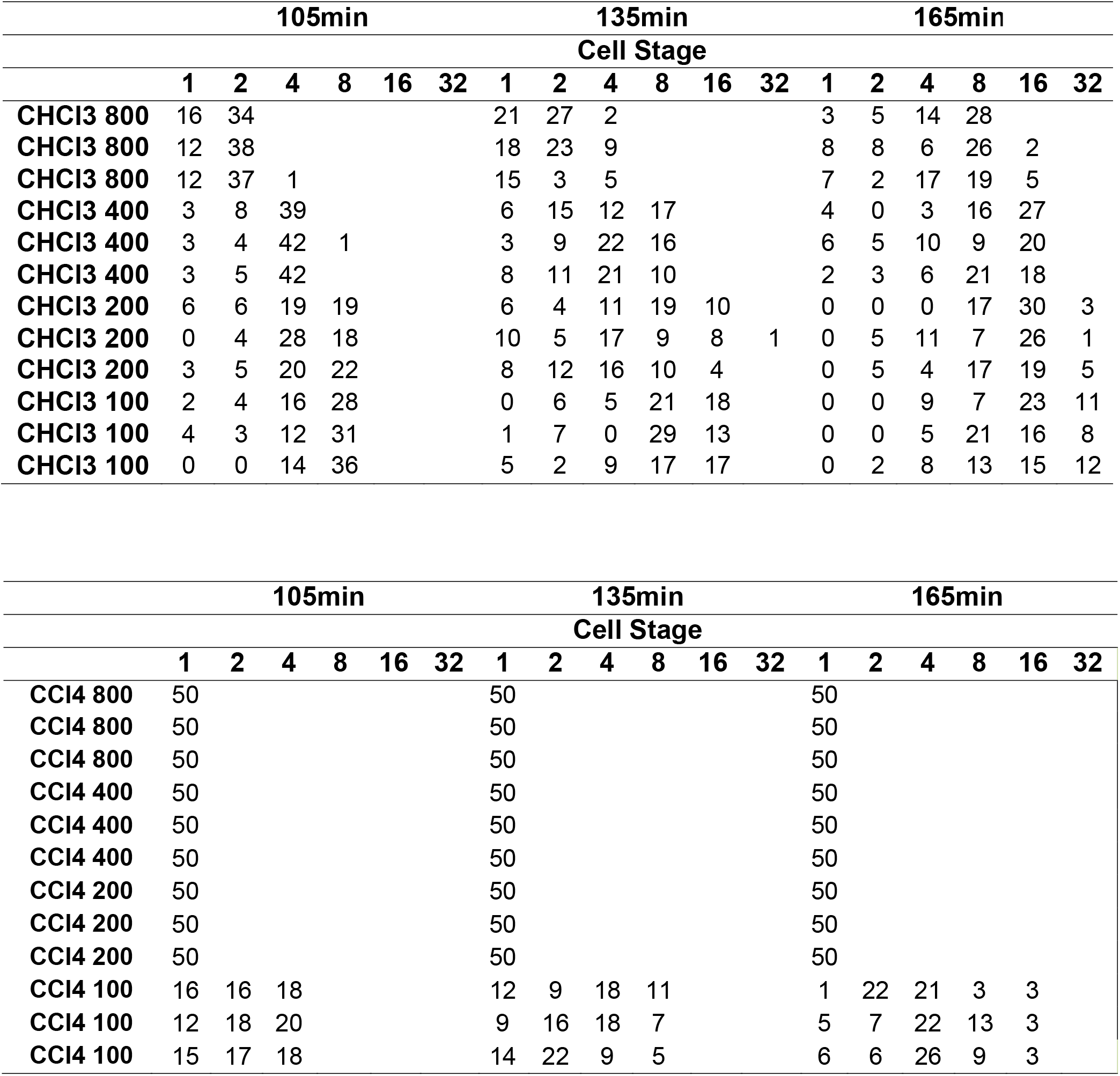
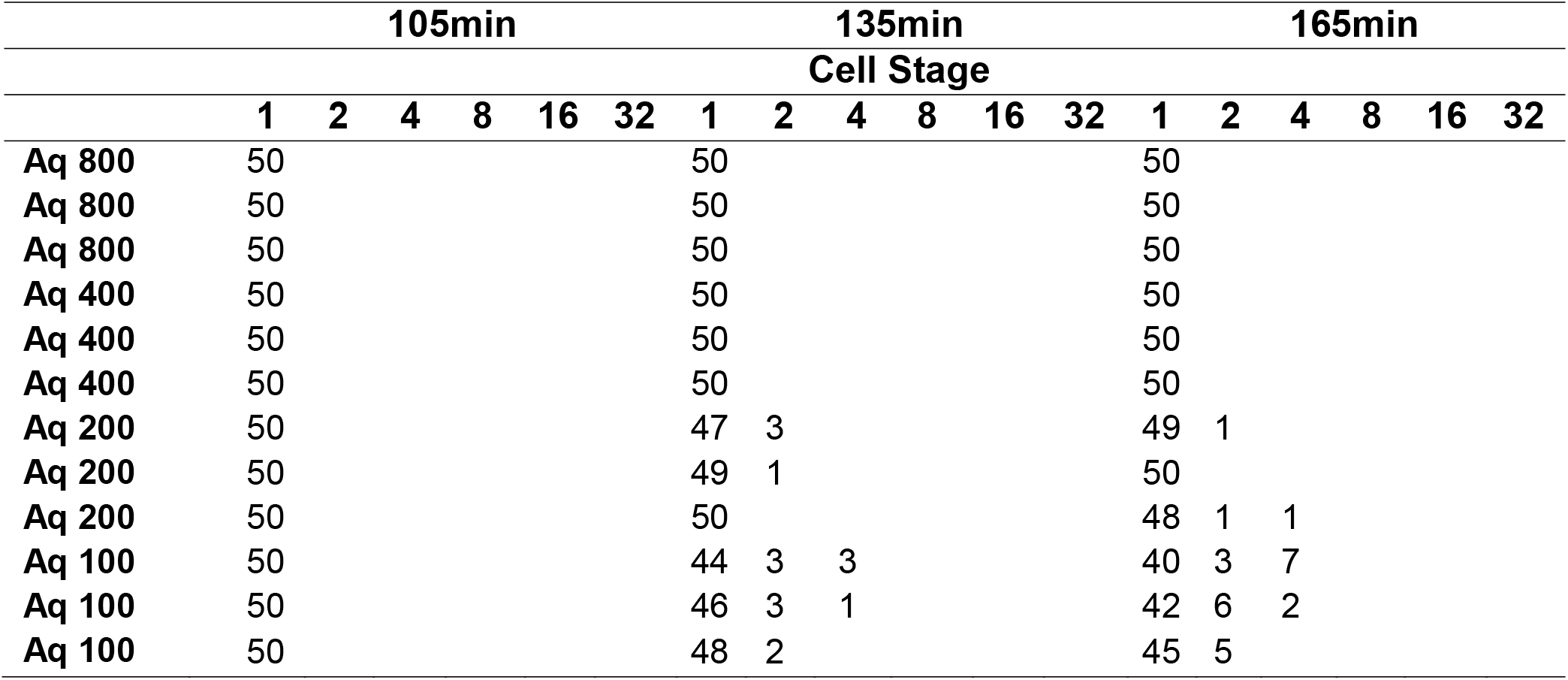

## APPENDIX D: Statistical Analysis

### I. ANOVA Raw Data

| B. Analysis of Variance (45min) |  |  |  |  |  |
| --- | --- | --- | --- | --- | --- |
| Source | DF | Adj SS | Adj MS | F - Value | P - Value |
| Factor | 17 | 0.004533 | 0.000267 | * | * |
| Error | 36 | 0 | 0 |  |  |
| Total | 53 | 0.00453 |  |  |  |

| Factor | N | Mean | St Dev |
| --- | --- | --- | --- |
| HEXIN800 | 3 | 1 | 0 |
| HEXIN400 | 3 | 1 | 0 |
| HEXIN200 | 3 | 1 | 0 |
| HEXIN100 | 3 | 1 | 0 |
| CHCL4800 | 3 | 1 | 0 |
| CHCL3400 | 3 | 1 | 0 |
| CHCL3200 | 3 | 1 | 0 |
| CHCL3100 | 3 | 1 | 0 |
| CCL4800 | 3 | 1 | 0 |
| CCL4400 | 3 | 1 | 0 |
| CCL4200 | 3 | 1 | 0 |
| CCL4100 | 3 | 1 | 0 |
| AQUO800 | 3 | 1 | 0 |
| AQUO400 | 3 | 1 | 0 |
| AQUO200 | 3 | 1 | 0 |
| AQUO100 | 3 | 1 | 0 |
| Positive | 3 | 1 | 0 |
| Negative | 3 | 1.04 | 0 |

Pooled St Dev = 0
| **C. Analysis of Variance (75min)** |  |  |  |  |  |
| --- | --- | --- | --- | --- | --- |
| **Source** | **DF** | **Adj SS** | **Adj MS** | **F - Value** | **P - Value** |
| **Factor** | 17 | 20.6843 | 1.21461 | 141.23 | 0 |
| **Error** | 36 | 0.3096 | 0.0086 |  |  |
| **Total** | 53 | 20.9579 |  |  |  |

| **Factor** | **N** | **Mean** | **St Dev** |
| --- | --- | --- | --- |
| HEXIN800 | 3 | 1 | 0 |
| HEXIN400 | 3 | 1 | 0 |
| HEXIN200 | 3 | 1 | 0 |
| HEXIN100 | 3 | 1.1 | 0.02 |
| CHCL3800 | 3 | 1.18 | 0.06 |
| CHCL3400 | 3 | 1.927 | 0.221 |
| CHCL3200 | 3 | 1.8 | 0.272 |
| CHCL3100 | 3 | 2.967 | 0 |
| CCL4800 | 3 | 1 | 0 |
| CCL4400 | 3 | 1 | 0 |
| CCL4200 | 3 | 1 | 0.1155 |
| CCL4100 | 3 | 1.2133 | 0 |
| AQUO800 | 3 | 1 | 0 |
| AQUO400 | 3 | 1 | 0 |
| AQUO200 | 3 | 1 | 0 |
| AQUO100 | 3 | 1 | 0 |
| Positive | 3 | 1 | 0 |
| Negative | 3 | 2.86 | 0 |
| **Pooled St Dev = 0.0927362** |  |  |  |

D. Analysis of Variance (105min)
| Source | DF | Adj SS | Adj MS | F - Value | P - Value |
| --- | --- | --- | --- | --- | --- |
| **Factor** | 17 | 194.611 | 11.4477 | 632.08 | 0 |
| **Error** | 36 | 0.652 | 0.0181 |  |  |
| **Total** | 53 | 195.263 |  |  |  |

| Factor | N | Mean | St Dev |
| --- | --- | --- | --- |
| HEXIN800 | 3 | 1 | 0 |
| HEXIN400 | 3 | 1 | 0 |
| HEXIN200 | 3 | 1 | 0 |
| HEXIN100 | 3 | 1.50667 | 0.01155 |
| CHCL3800 | 3 | 1.7467 | 0.0611 |
| CHCL3400 | 3 | 3.62 | 0.12 |
| CHCL3200 | 3 | 5.193 | 0.242 |
| CHCL3100 | 3 | 6.32 | 0.492 |
| CCL4800 | 3 | 1 | 0 |
| CCL4400 | 3 | 1 | 0 |
| CCL4200 | 3 | 1 | 0 |
| CCL4100 | 3 | 2.46 | 0.0872 |
| AQUO800 | 3 | 1 | 0 |
| AQUO400 | 3 | 1 | 0 |
| AQUO200 | 3 | 1 | 0 |
| AQUO100 | 3 | 1 | 0 |
| Positive | 3 | 1 | 0 |
| Negative | 3 | 6.72 | 0 |

**Pooled St Dev = 0.134578** E. Analysis of Variance (135min)
| Source | DF | Adj SS | Adj MS | F - Value | P - Value |
| --- | --- | --- | --- | --- | --- |
| **Factor** | 17 | 818.618 | 48.154 | 513.37 | 0 |
| **Error** | 36 | 30377 | 0.0938 |  |  |
| **Total** | 53 | 821.995 |  |  |  |

| Factor | N | Mean | St Dev |
| --- | --- | --- | --- |
| **HEXIN800** | 3 | 1 | 0 |
| **HEXIN400** | 3 | 1 | 0 |
| **HEXIN200** | 3 | 1.02 | 0.0346 |
| **HEXIN100** | 3 | 1.3933 | 0.1102 |
| **CHCL3800** | 3 | 1.853 | 0.175 |
| **CHCL3400** | 3 | 4.34 | 0.433 |
| **CHCL3200** | 3 | 7.32 | 0.985 |
| **CHCL3100** | 3 | 9.307 | 0.393 |
| **CCL4800** | 3 | 1 | 0 |
| **CCL4400** | 3 | 1 | 0 |
| **CCL4200** | 3 | 1 | 0 |
| **CCL4100** | 3 | 3.287 | 0.566 |
| **AQUO800** | 3 | 1 | 0 |
| **AQUO400** | 3 | 1 | 0 |
| **AQUO200** | 3 | 1.0267 | 0.0306 |
| **AQUO100** | 3 | 1.1333 | 0.1007 |
| **Positive** | 3 | 1 | 0 |
| **Negative** | 3 | 15.86 | 0 |
| **Pooled St Dev = 0.306268** |  |  |  |

| **F. Analysis of Variance (165min)** |  |  |  |  |  |
| --- | --- | --- | --- | --- | --- |
| **Source** | **DF** | **Adj SS** | **Adj SS2** | **F - Value** | **P - Value** |
| **Factor** | 17 | 2922.77 | 171.927 | 522.65 | 0 |
| **Error** | 36 | 11.84 | 0.329 |  |  |
| **Total** | 53 | 2934.61 |  |  |  |

| **Factor** | **N** | **Mean** | **St Dev** |
| --- | --- | --- | --- |
| **HEXIN800** | 3 | 1.000 | 0.000 |
| **HEXIN400** | 3 | 1.000 | 0.000 |
| **HEXIN200** | 3 | 1.2533 | 0.0833 |
| **HEXIN100** | 3 | 3.687 | 0.318 |
| **CHCL3800** | 3 | 5.947 | 0.242 |
| **CHCL3400** | 3 | 10.080 | 0.310 |
| **CHCL3200** | 3 | 12.640 | 1.544 |
| **CHCL3100** | 3 | 15.173 | 1.124 |
| **CCL4800** | 3 | 1.000 | 0.000 |
| **CCL4400** | 3 | 1.000 | 0.000 |
| **CCL4200** | 3 | 1.000 | 0.000 |
| **CCL4100** | 3 | 4.680 | 0.596 |
| **AQUO800** | 3 | 1.000 | 0.000 |
| **AQUO400** | 3 | 1.000 | 0.000 |
| **AQUO200** | 3 | 1.0333 | 0.0416 |
| **AQUO100** | 3 | 1.273 | 0.192 |
| **Pos control** | 3 | 1.020 | 0.000 |
| **Neg control** | 3 | 29.76 | 0.00 |

### II. Post-hoc Tukey Test Data

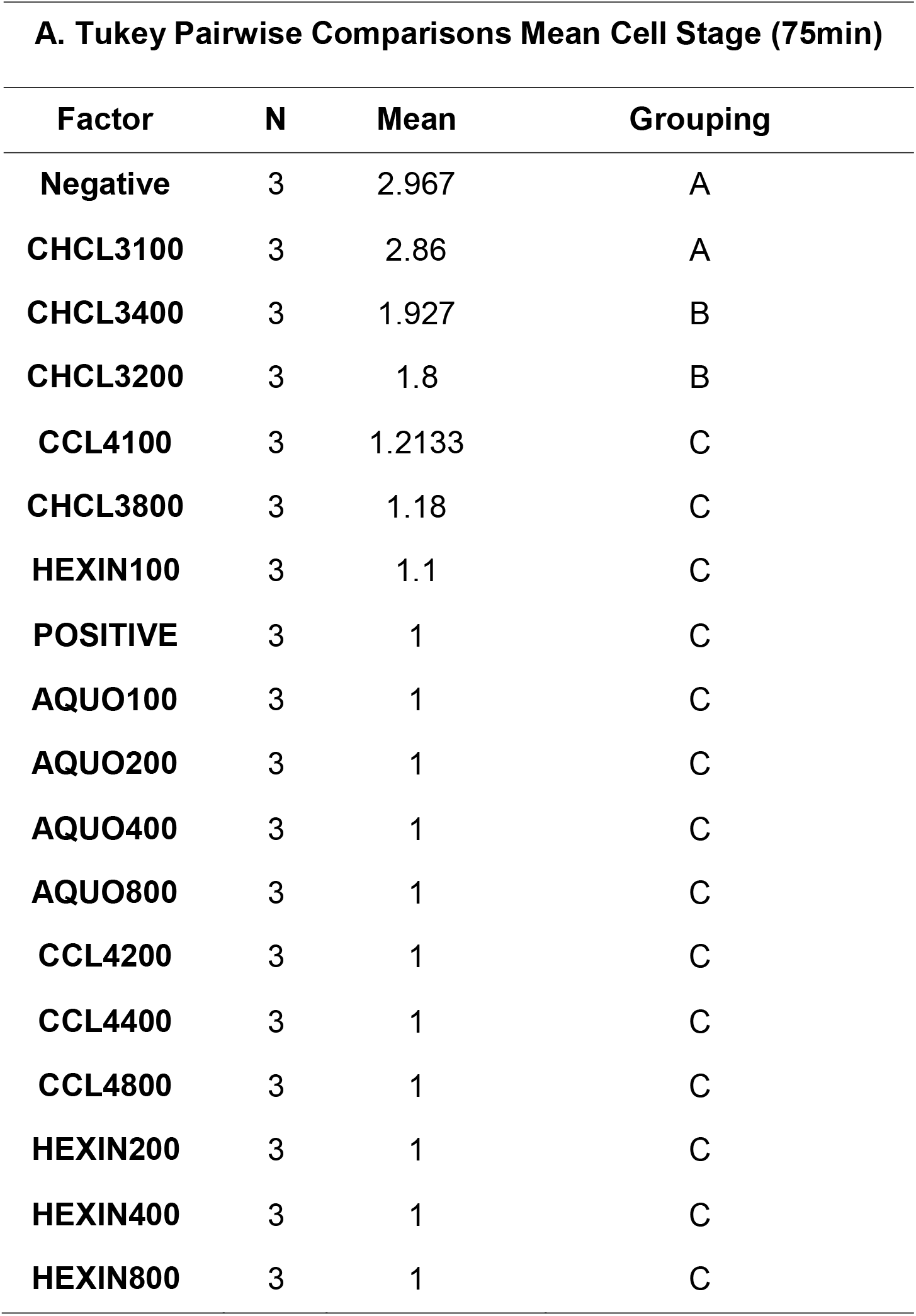
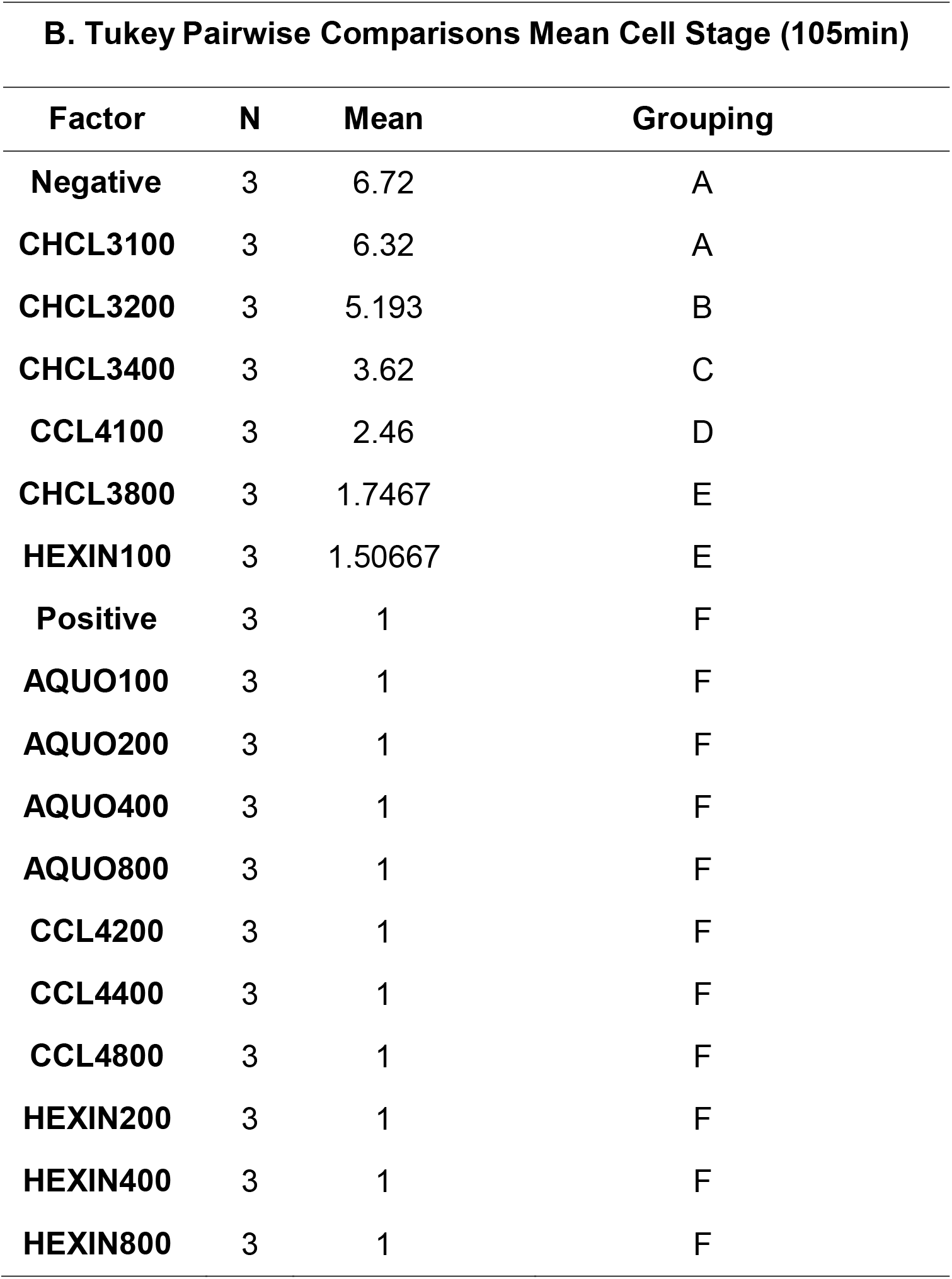
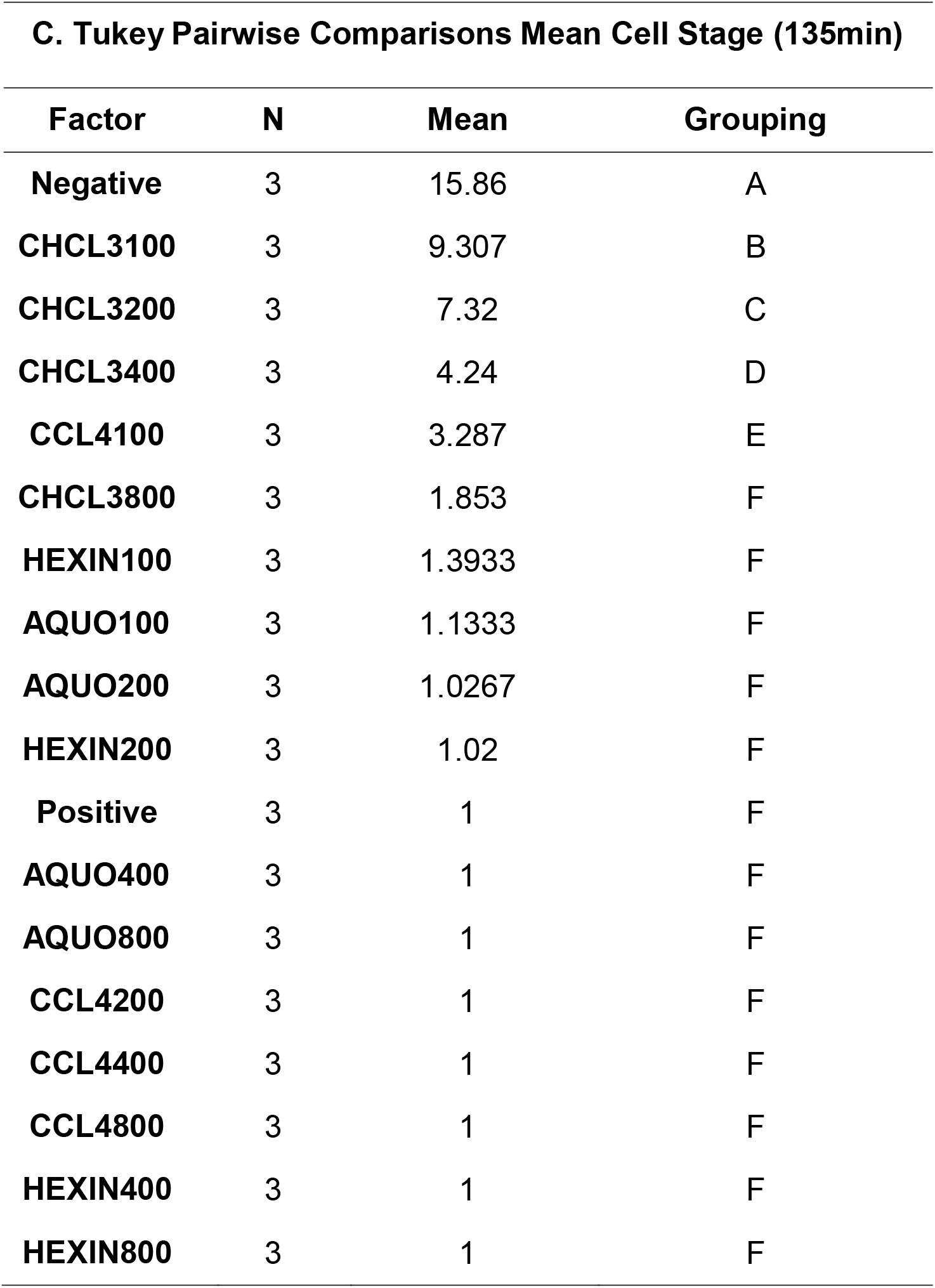
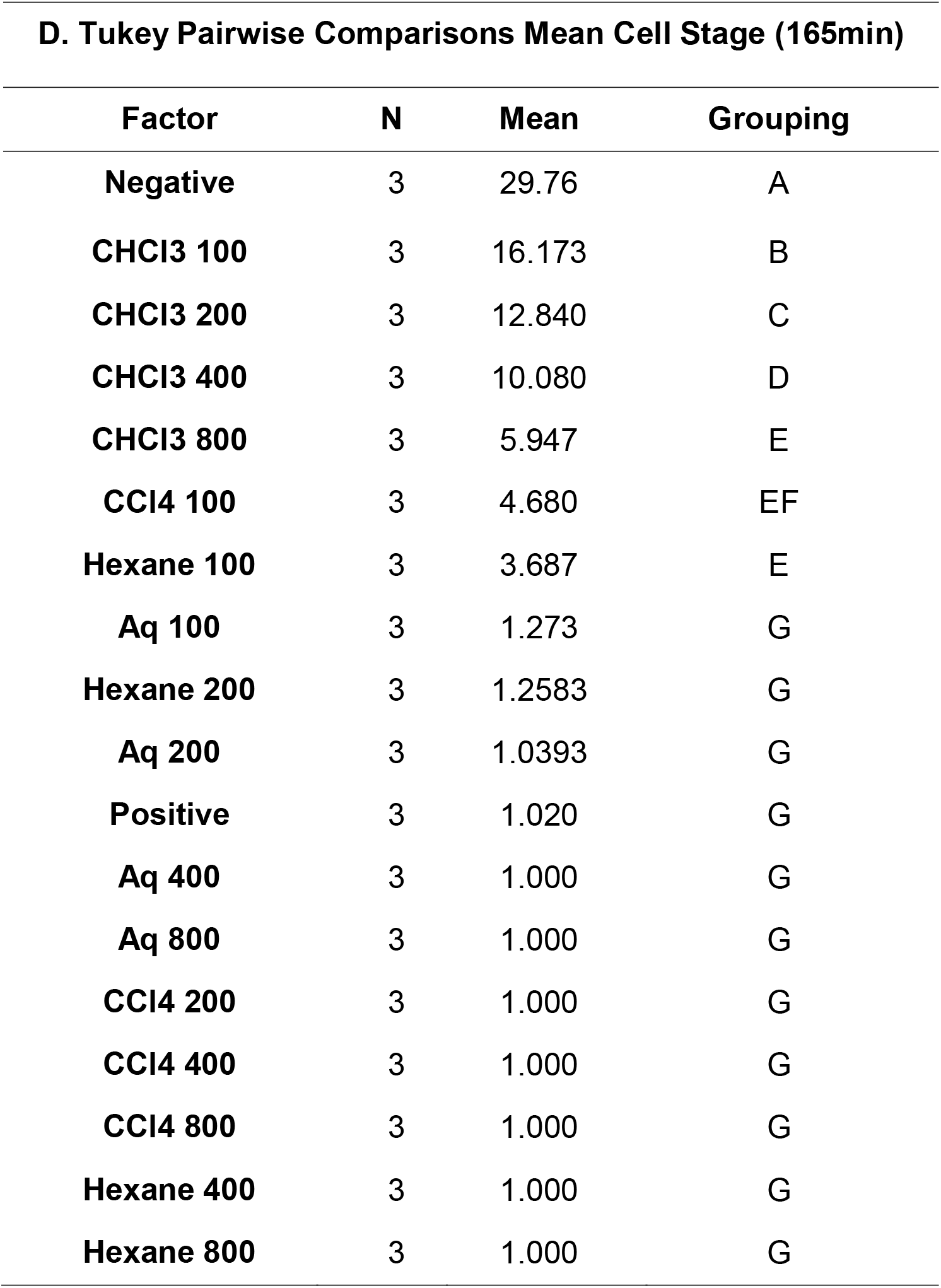

## APPENDIX E: Pictures

### I. *Samanea saman* leaves collection

**Fig 1.**
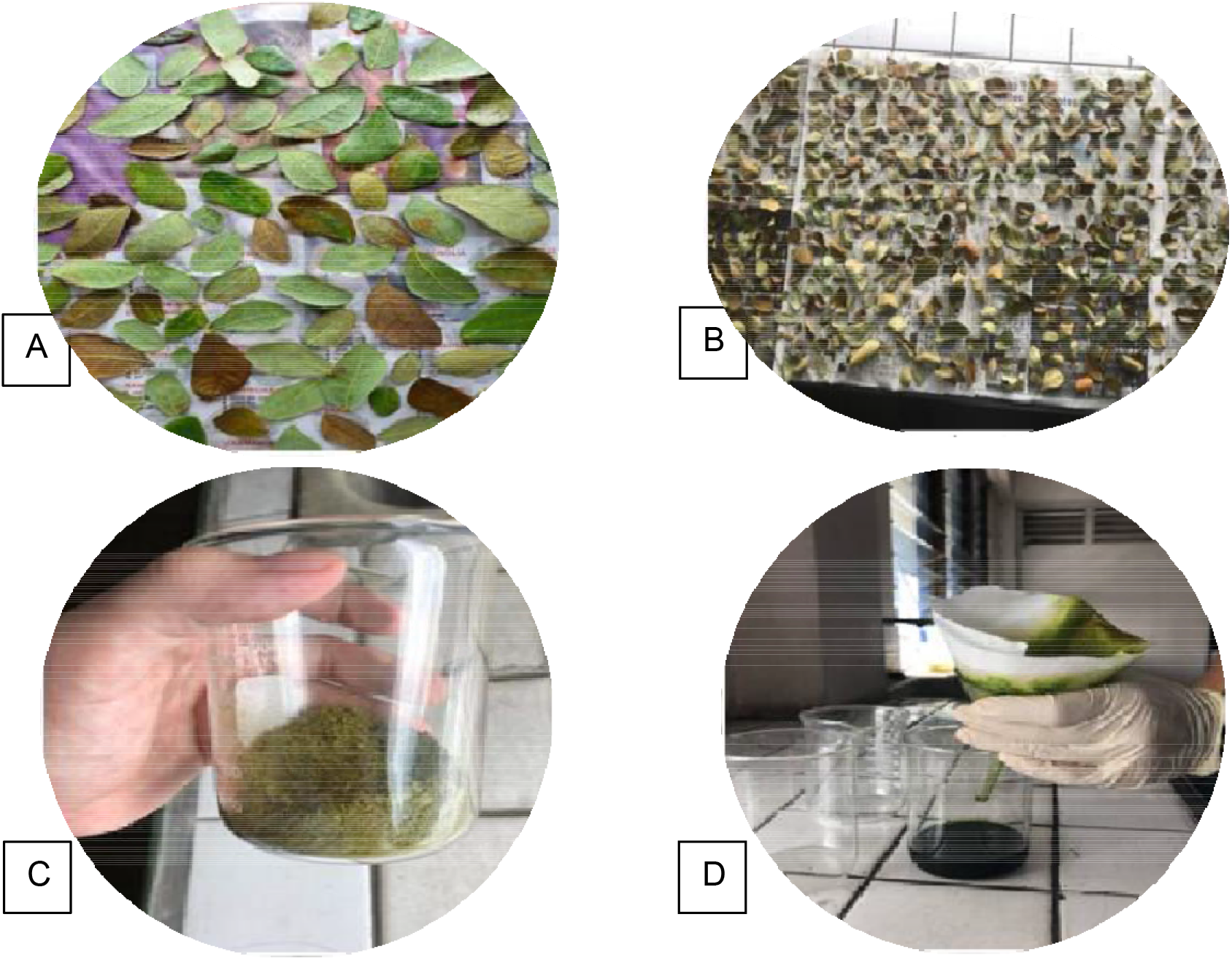
*S. saman* leaf extraction process: (A) Fresh *S. saman* leaves, (B) Dried *S. saman* leaves, (C) Pulverized dried *S. saman* leaves, (D) Maceration and filtration of *S. saman* extract

### II. *Samanea saman* leaves extraction

**Fig. 2.**
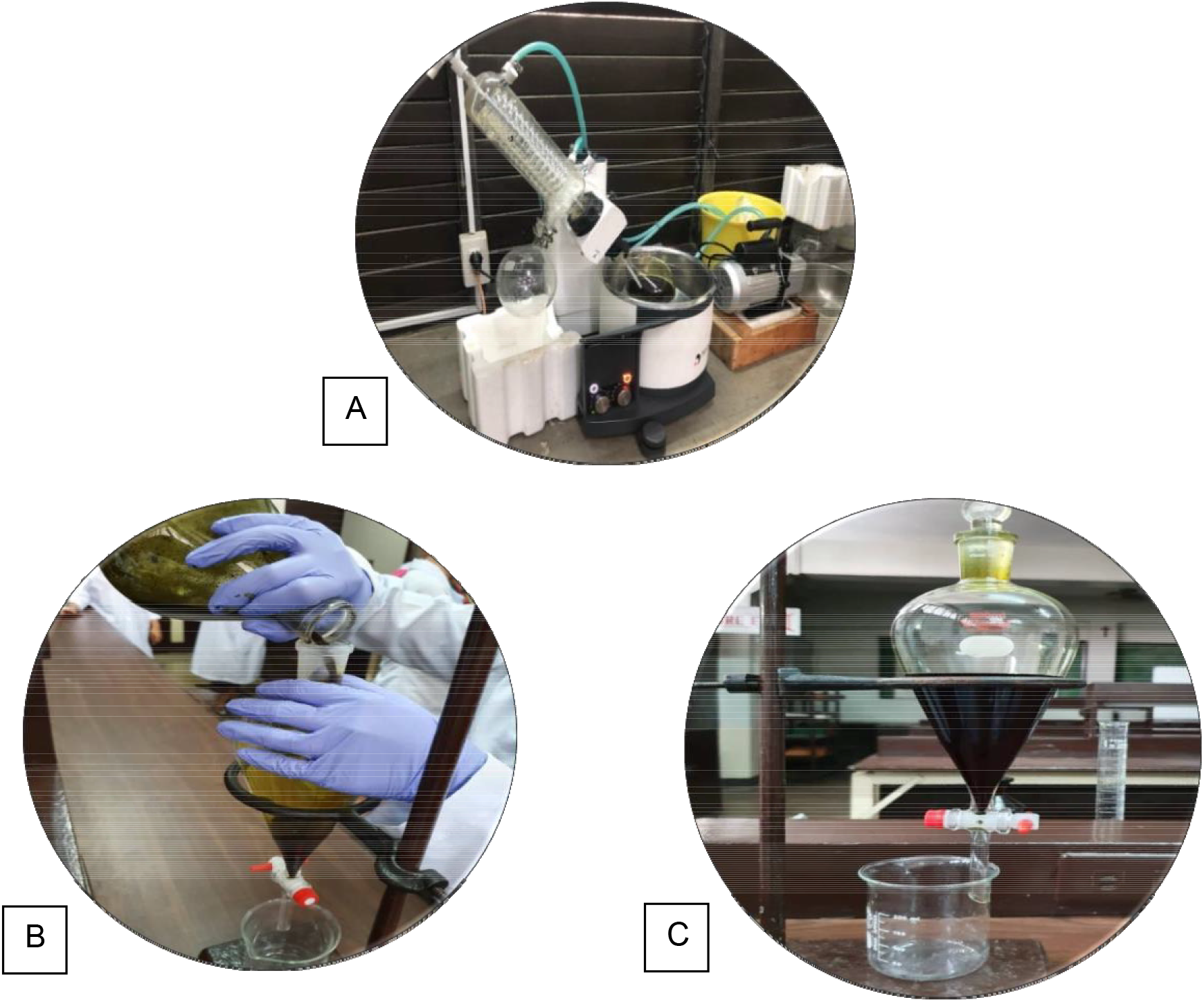
*S. saman* extraction: (A) Rotary evaporation of macerated *S. saman* leaves, (B) Pouring of extract into separatory funnel, (C) Separation of polar and nonpolar compounds

### III. *Tripneustes gratilla* sperm and egg spawning

**Fig 3.**
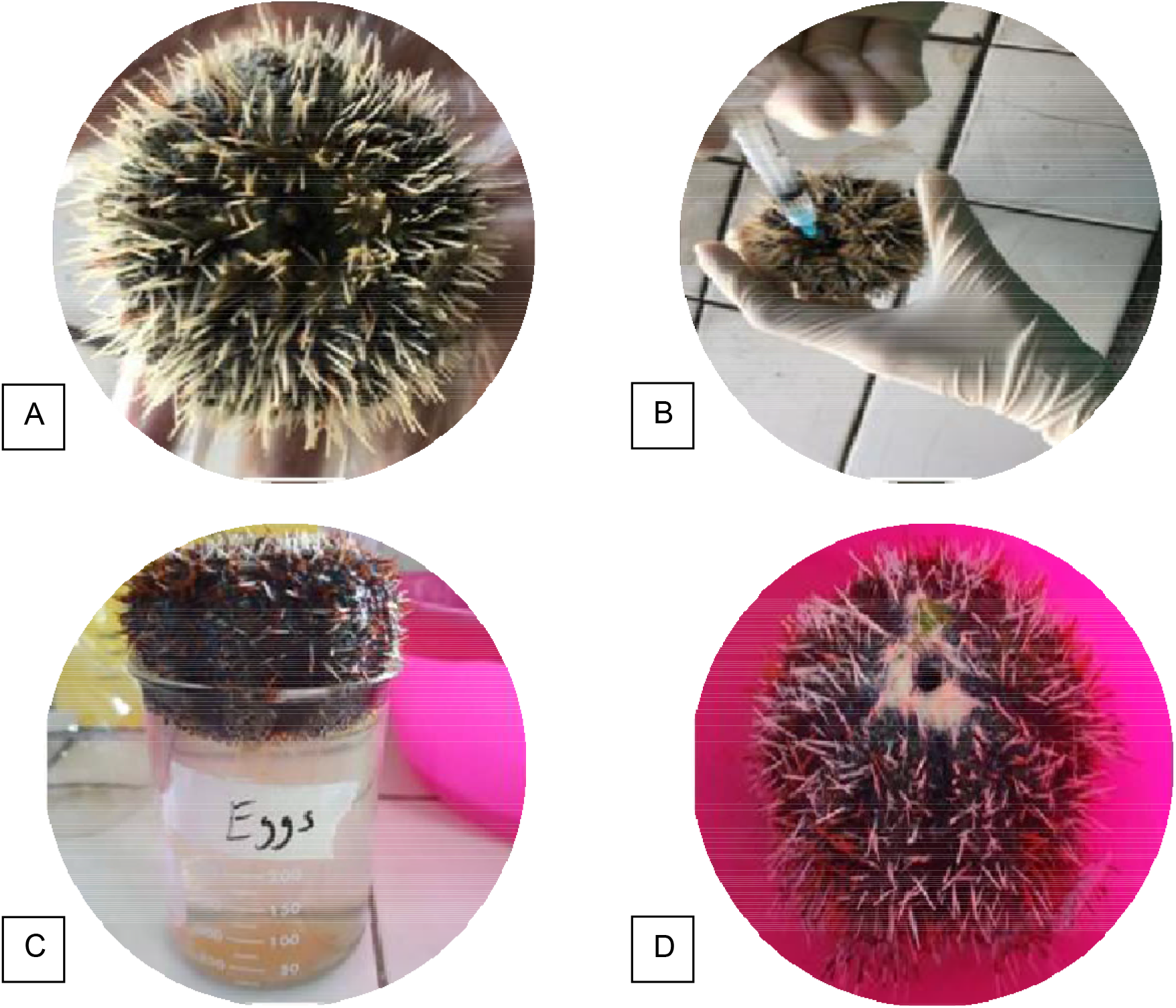
*T. gratilla* gametes production: (A) *T. gratilla* test subject, (B) 1% KOH injected at mouth side up of sea urchin, (C) Female *T. gratilla* spawning eggs, (D) Male *T. gratilla* spawning sperms

### IV. Cell division stages of *Tripneustes gratilla*

**Fig. 4.**
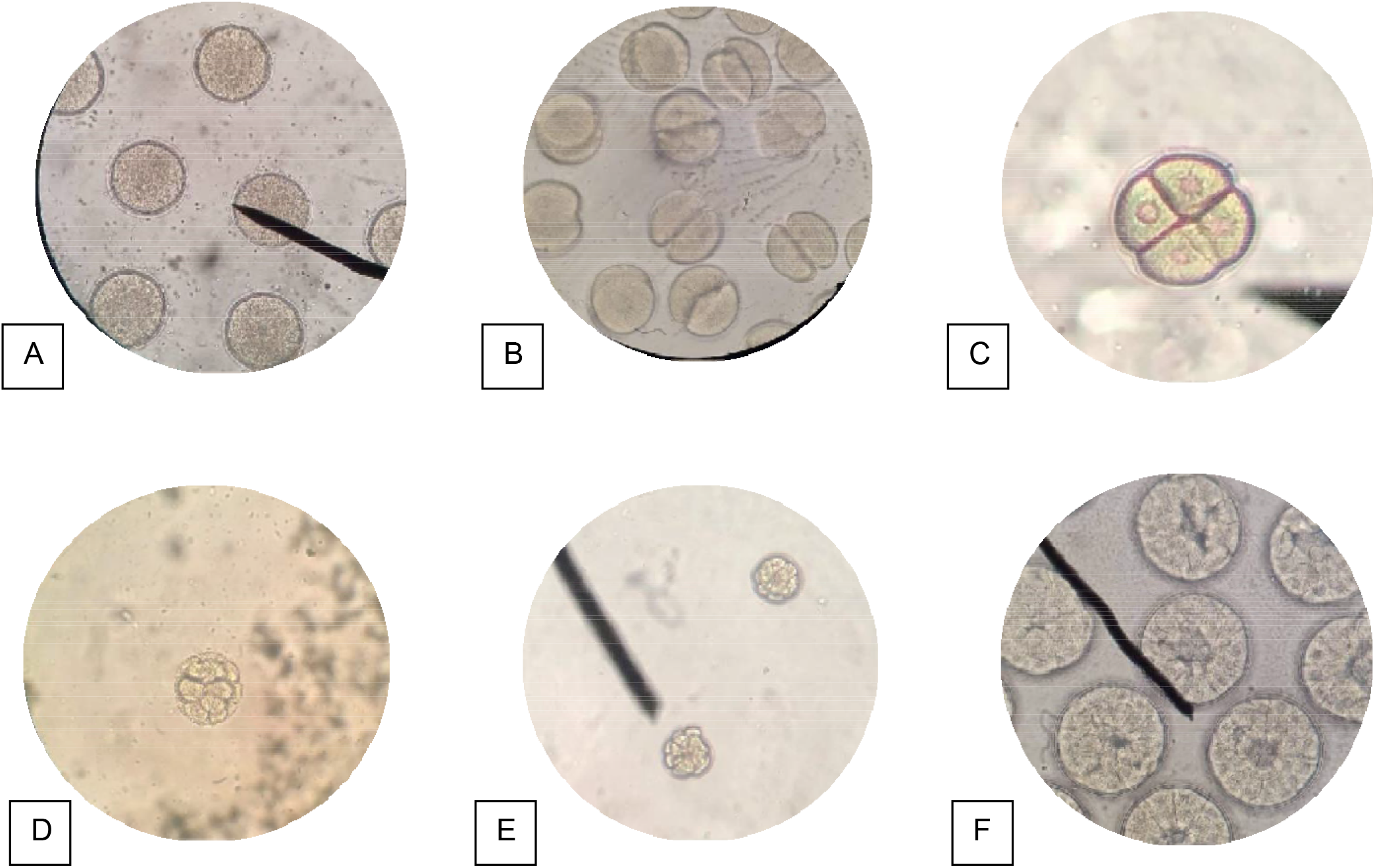
Cell division stages of *T. gratilla*: (A) Fertilization envelope, (B) 2-cell stage, (C) 4-cell stage, (D) 8-cell stage, (E) 16-cell stage, (F) 32-cell stage

### V. Disposal of used *Tripneustes gratilla*

**Fig 5.**
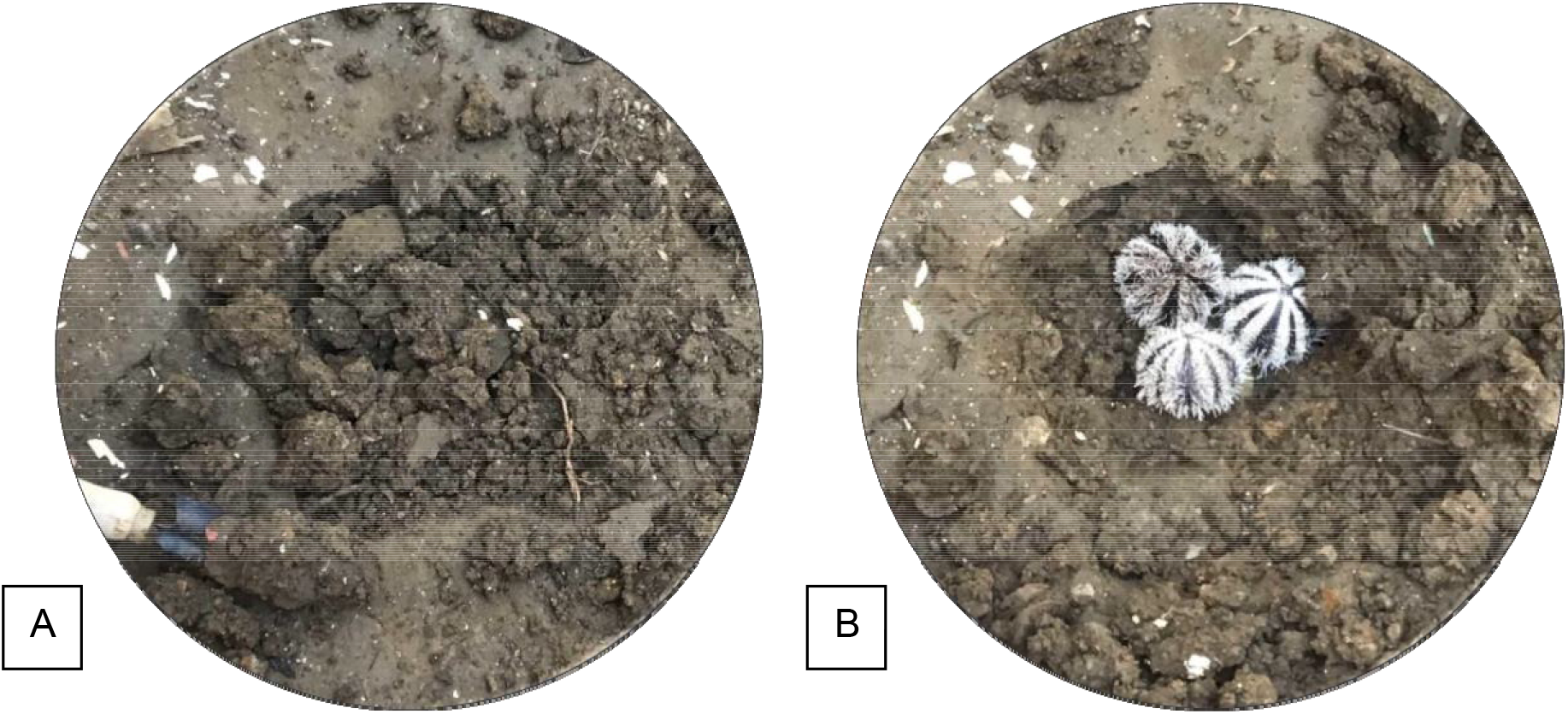
(A, B) Euthanized *T. gratilla* disposal

## APPENDIX F: Financial Table

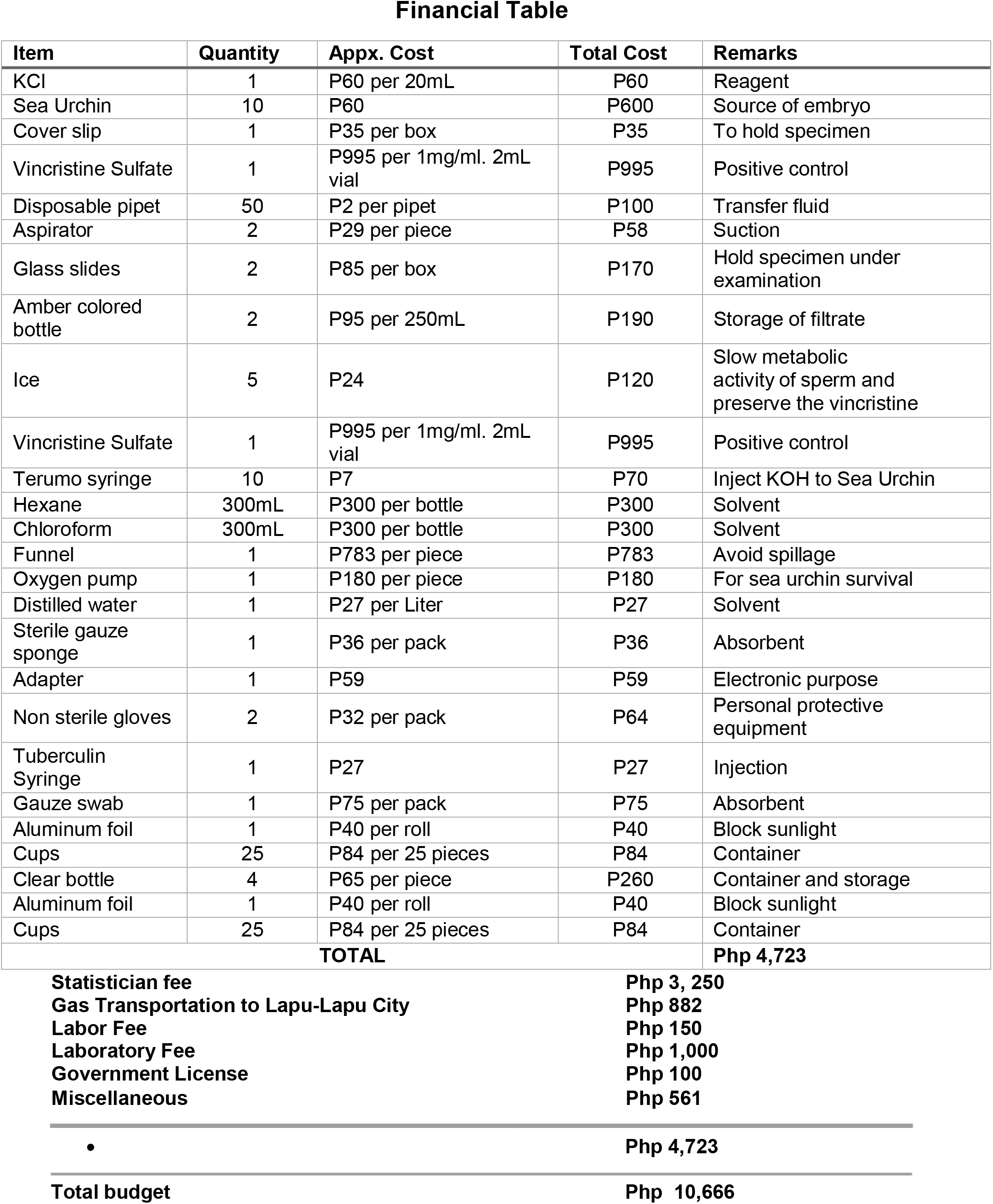

## APPENDIX G: Gantt Chart

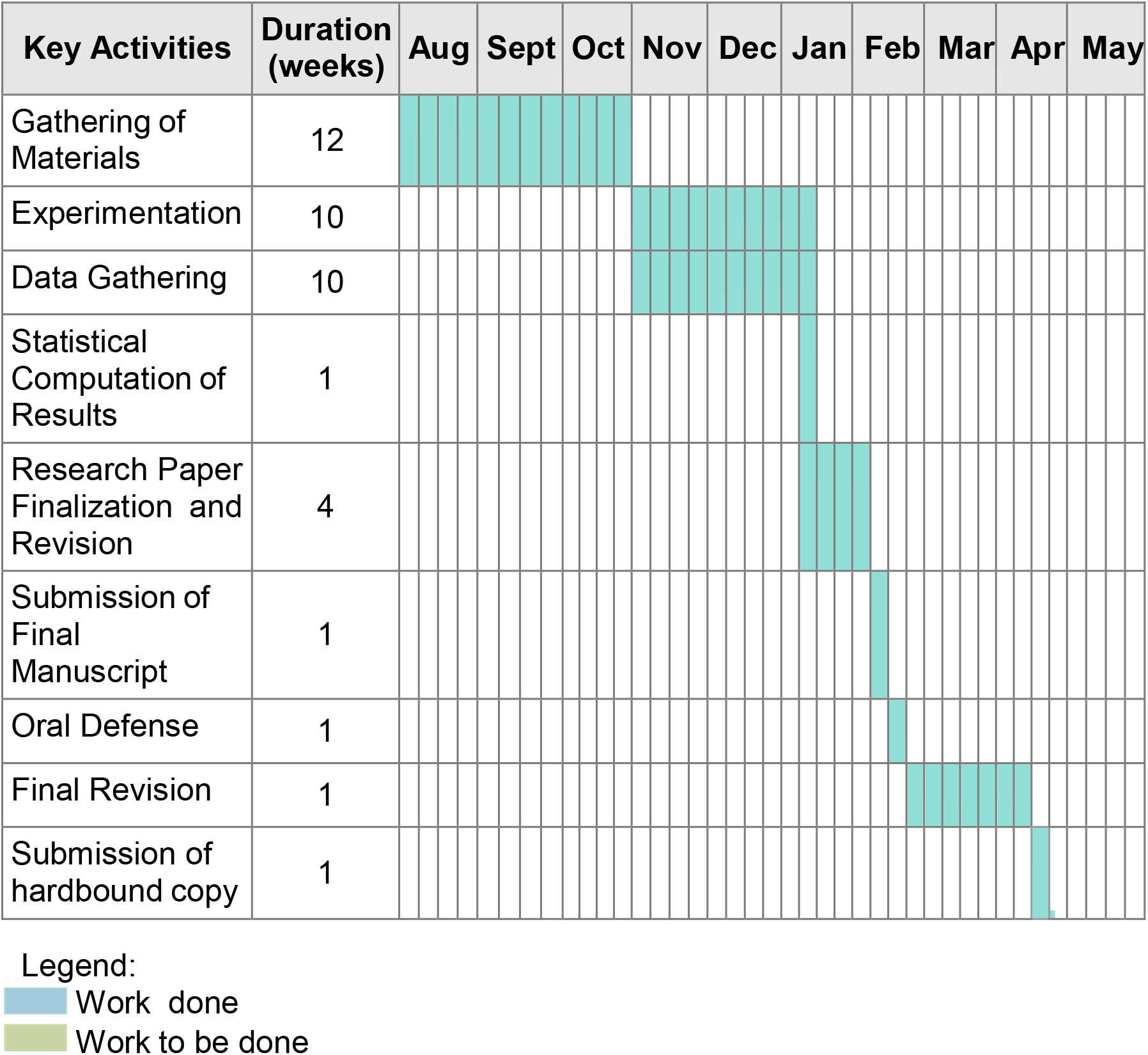

